# A tumor–pituitary–stromal axis promotes colorectal cancer progression

**DOI:** 10.64898/2026.09.16.751939

**Authors:** Weihan Li, Shibo Wang, Hao Wu, Chenjie Shen, Sailiang Liu, Hao Wang, Ruixue Huo, Kexin He, Chuhan Zhang, Minhao Yu, Shu-Heng Jiang, Junli Xue

## Abstract

Peripheral tumors can reprogram systemic physiology, yet how they engage central endocrine organs to remodel the local tumor microenvironment is unknown. We found that colorectal cancer (CRC) elevates circulating BMP2, which activates pituitary signaling and enhances endocrine output. Increased pituitary-derived growth hormone acted on growth hormone receptor–expressing cancer-associated fibroblasts (CAFs), inducing a p21-dependent senescence-like state and tumor-promoting secretory program. Disruption of pituitary BMP signaling or fibroblast GHR reduced tumor growth, whereas fibroblast-specific depletion of p21 abrogated GHR-dependent tumor promotion. In patients with CRC, elevated pituitary metabolic activity correlated with stromal p21 expression and tumor burden. Our study reveals a tumor–pituitary–stromal signaling axis through which a peripheral malignancy recruits a central endocrine organ to remotely shape its own microenvironment and promote disease progression.

## Main Text

Cancer is increasingly recognized as a systemic disease in which tumor progression is shaped by reciprocal interactions between malignant tissues and the host (*1–3*). Beyond the local tumor microenvironment (TME), tumor-derived signals can enter the circulation and remodel distant tissues and physiological regulatory systems, including immune, metabolic, neural, and endocrine networks. These systemic responses are not necessarily passive consequences of disease but can feed back on tumors to influence progression, treatment response, and other cancer-associated phenotypes. Tumors can also disrupt systemic endocrine homeostasis and exploit circulating hormonal signals to support disease progression (*4*). As a central endocrine organ, the pituitary is well positioned to mediate such long-range tumor–host communication by integrating systemic inputs and coordinating diverse physiological processes through hormone secretion (*5–7*). Previous studies have shown that pituitary-derived hormones can influence tumor growth, immune regulation, and therapeutic response (*8–10*), highlighting the potential importance of pituitary endocrine activity in cancer. However, whether a peripheral tumor actively remodels the pituitary, how this endocrine response is induced, and whether it functionally contributes to tumor development remain largely unknown.

Cancer-associated fibroblasts (CAFs) are highly plastic components of the TME that adopt diverse functional states in response to cytokines, growth factors, metabolic cues, and tissue stress, thereby regulating tumor growth, immune evasion, extracellular matrix remodeling, and therapeutic response (*11, 12*). Senescence-associated programs can further endow fibroblasts with persistent tumor-promoting properties (*13, 14*). Although CAF states are generally viewed as products of local tumor-derived signals, whether systemic endocrine cues contribute to their functional remodeling remains unclear. Here, we investigated whether colorectal cancer (CRC) engages the pituitary as an intermediate endocrine node to regulate the tumor stroma. Through region-resolved transcriptomic profiling, pituitary single-nucleus sequencing, plasma proteomics, genetic mouse models, functional perturbation, and analyses of patients with CRC, we identify a tumor–pituitary–stromal axis in which CRC-associated systemic signals activate pituitary endocrine output and remodel CAF states to support tumor progression.

### CRC induces transcriptomic remodeling and promotes hormone secretion in the pituitary gland

To investigate the effects of CRC on the brain and pituitary gland, mice were euthanized 28 days after orthotopic tumor induction via submucosal injection of MC38 cells into the rectal wall **(fig. S1, A to C)** (*15*). The brain was then dissected into five defined regions, including the cerebellum, hindbrain, hypothalamus, thalamus with midbrain, and cerebrum, together with the pituitary gland, for region-specific RNA sequencing (RNA-seq) **(Fig. 1A)**. When we compared transcriptional profiles across five brain regions and the pituitary between control and MC38 tumor-bearing mice, principal component analysis (PCA) showed clear segregation by anatomical region. Of note, pituitary samples showed a clear divergence between PBS and MC38 groups along PC1, which explains the most variance in the data **(Fig. 1, B and C)**. Gene expression analysis further revealed that the pituitary harbored the largest number of differentially expressed genes (DEGs) among the analyzed regions **(fig. S1D)**.

**Fig. 1.**
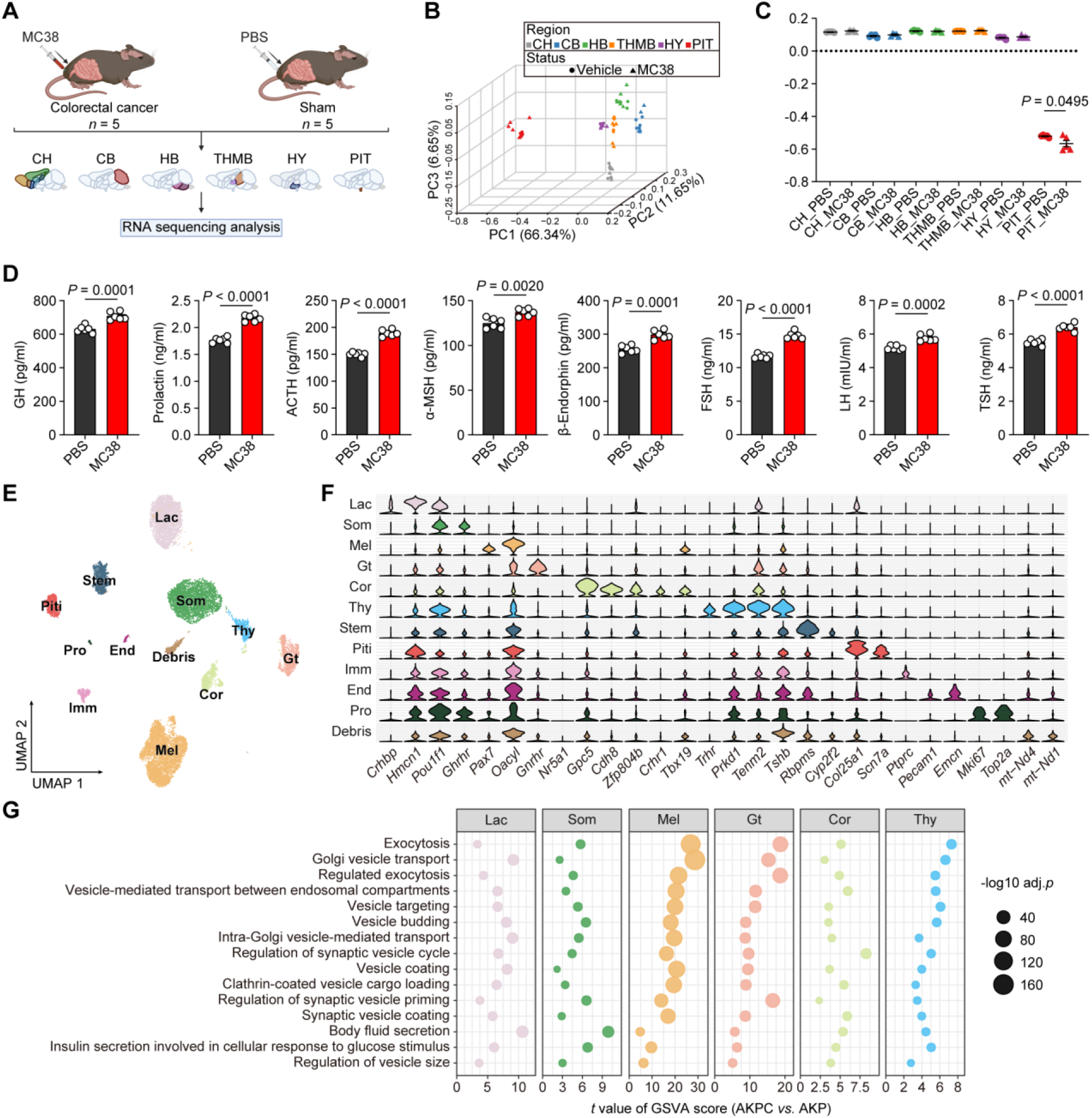
CRC induces pituitary transcriptomic remodeling and enhances hormone secretion. (**A** to **C**) Bulk RNA-seq analysis of dissected brain regions and pituitary gland. (A) RNA-seq profiling summary. An orthotopic CRC model was established by submucosal injection of MC38 cells into the rectum of female C57BL/6J mice (*n* = 5). Sham mice received an equal-volume PBS injection (*n* = 5). After tumor formation, distinct brain regions were dissected for RNA-seq analysis. CH, cerebrum; CB, cerebellum; HB, hindbrain; THMB, thalamus with midbrain; HY, hypothalamus; PIT, pituitary. (B) PCA analysis. (C) Quantification of PC1 scores in (B). (**D**) ELISA assays of plasma adenohypophyseal hormones of control and MC38 tumor-bearing mice at 28 d after MC38 tumor implantation (*n* = 5 mice/group). GH, growth hormone; ACTH, adrenocorticotropic hormone; α-MSH, α-melanocyte-stimulating hormone; FSH, follicle-stimulating hormone; LH, luteinizing hormone; TSH, thyroid-stimulating hormone. (**E** to **G**) snRNA-seq analysis of pituitary. (E) Uniform manifold approximation and projection (UMAP) of cell clusters. (F) Violin plot of selected cluster-specific genes. (G) Differential pathway enrichment in pituitary endocrine cell populations by GSVA. Two-tailed unpaired limma-moderated *t* test. The data are presented as mean ± SEM. *P* values are determined by two-tailed unpaired Student’s *t* test (C and D). Figure A created with BioRender.com.

To validate the spatial specificity of our dataset, we examined hormone-related gene expression. Neurohypophysial genes (*Avp, Oxt*) were enriched in the hypothalamus, whereas adenohypophyseal genes (*Gh, Prl, Pomc, Cga, Fshb, Lhb,* and *Tshb*) were enriched in the pituitary **(fig. S1E)**. We therefore focused on adenohypophyseal hormones to assess CRC-induced alterations in pituitary endocrine function. Notably, plasma levels of adenohypophyseal hormones were increased in MC38 tumor-bearing mice, including GH, prolactin, adrenocorticotropic hormone (ACTH), α-melanocyte-stimulating hormone (α-MSH), β-endorphin, follicle-stimulating hormone (FSH), luteinizing hormone (LH), and thyroid-stimulating hormone (TSH) **(Fig. 1D)**.

We next examined the AKPC autochthonous CRC mouse model, which recapitulates the progression of human CRC from adenoma to adenocarcinoma and invasive disease **(fig. S2, A to C)**. AKPC mice exhibited elevated plasma levels of multiple adenohypophyseal hormones compared with littermate AKP controls (*Apc^15lox/+^;Kras^LSL-G12D/+^;Trp53^R172H/+^*), with significant increases in GH, prolactin, ACTH, α-MSH, LH, and TSH **(fig. S2D)**.

To further understand how CRC influences the pituitary, we performed single-nucleus RNA sequencing (snRNA-seq) of pituitary glands isolated from AKP and AKPC mice. We identified eleven distinct cell populations, including six endocrine cell subsets (lactotropes, somatotropes, melanotropes, gonadotropes, corticotropes, and thyrotropes), as well as stem cells, pituicytes, immune cells, endothelial cells, and proliferating cells **(Fig. 1E)**. Cell identities were annotated and consolidated based on marker genes as described by Ruf-Zamojski *et al.* **(Fig. 1F)** (*16*).

These annotations were further supported by pathway activity analysis among pituitary cell populations **(fig. S2E)**. Further evaluation of pathway activity in pituitary endocrine populations using gene set variation analysis (GSVA) revealed a shared increase in exocytosis- and vesicle trafficking–related pathways, consistent with enhanced hormone secretion in AKPC mice compared with AKP controls **(Fig. 1G)**. These results suggest that, despite their functional diversity, pituitary endocrine cell types exhibit an overlapping pattern of responses to CRC.

### CRC triggers pituitary responses via BMP2

Tumors can modulate pituitary function through circulating mediators (*4*). To explore factors that regulate pituitary function, we profiled the plasma proteome of control and MC38 tumor-bearing mice **(table S1)**. A total of 208 proteins were upregulated and 165 were downregulated in tumor-bearing mice compared with controls **(fig. S3A)**. Gene ontology (GO) enrichment analysis revealed that the significantly regulated proteins were mainly enriched in the “extracellular space” and “extracellular region,” suggesting a predominance of secreted proteins **(fig. S3B)**. We next analyzed proteins annotated as secreted in the UniProt database (Swiss-Prot, subcellular location; downloaded April 2025 (*17*)) **(table S2)**. Among the 208 upregulated proteins in MC38 tumor-bearing mice, 26 were annotated as secreted according to UniProt subcellular localization **(fig. S3C)**, five of which have documented receptor(s) based on ligand–receptor analysis using CellPhoneDB **(Fig. 2A)**. We further analyzed the expression of these receptors using bulk RNA-seq and snRNA-seq datasets of the mouse pituitary **(Fig. 2, B and C, and fig. S3D)**. Through this screening, bone morphogenetic protein 2 (BMP2) was identified as a candidate protein for in-depth functional investigation.

**Fig. 2.**
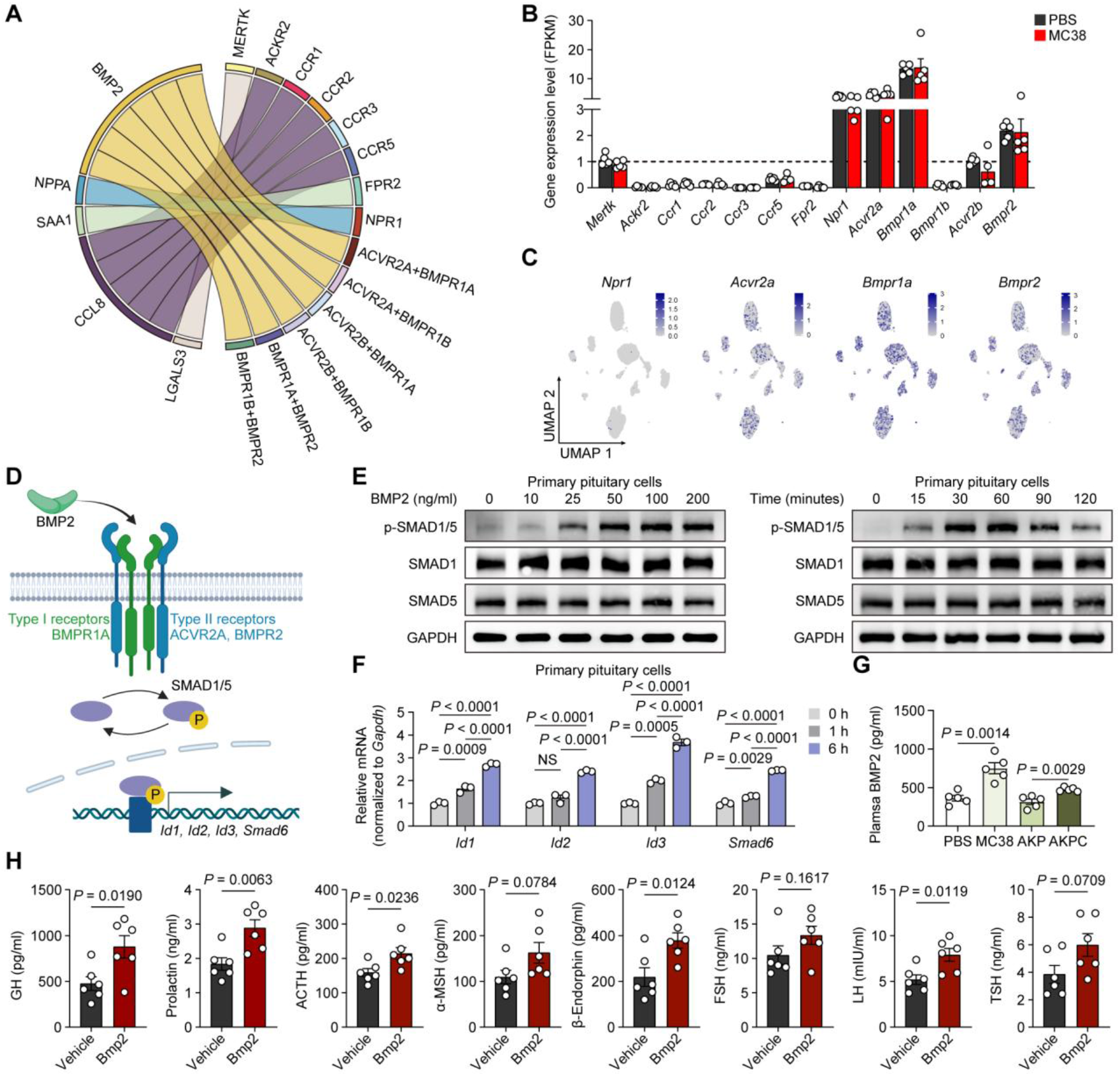
CRC triggers pituitary responses via BMP2. (**A**) Ligand–receptor analysis of upregulated secreted plasma proteins in MC38 tumor-bearing mice. (**B**) Bulk RNA-seq analysis showing the expression of receptors corresponding to the candidate secreted proteins in mouse pituitary tissues from PBS control and MC38 tumor-bearing mice. The dashed line indicates an FPKM value of 1. (**C**) snRNA-seq UMAP feature plots showing the pituitary distribution of receptors for candidate proteins prioritized from the bulk RNA-seq analysis in (B). (**D**) Illustration of BMP pathway components. (**E**) Immunoblot analysis of primary pituitary cells treated with recombinant mouse BMP2 (rmBMP2) at the indicated concentrations for 45 min (left) or with 25 ng/mL rmBMP2 for the indicated durations (right). Glyceraldehyde-3-phosphate dehydrogenase (GAPDH) was used as a loading control. (**F**) qRT-PCR analysis of the mRNA expression of the SMAD1/5 target genes *Id1, Id2, Id3 and Smad6* in primary pituitary cells. Fold change (FC) relative to unstimulated cells is shown for 1 h and 6 h stimulation with 25 ng/mL rmBMP2 (*n* = 3 biological replicates). (**G**) BMP2 ELISA of plasma samples from MC38 orthotopic tumor-bearing mice and AKPC spontaneous tumor-bearing mice (*n* = 5 mice per group). (**H**) ELISA analysis of plasma pituitary hormones in non-tumor-bearing C57BL/6J wild-type mice after intraperitoneal administration of recombinant mouse BMP2 (rmBMP2) for three consecutive days (*n* = 6 mice per group). GH, growth hormone; PRL, prolactin; ACTH, adrenocorticotropic hormone; α-MSH, α-melanocyte-stimulating hormone; FSH, follicle-stimulating hormone; LH, luteinizing hormone; TSH, thyroid-stimulating hormone. The data are presented as mean ± SEM. *P* values are determined by one-way analysis of variance (ANOVA) with Tukey’s multiple comparisons test (F) and two-tailed unpaired Student’s *t* test (G and H). NS, not significant. Figure D created with BioRender.com.

As a developmental morphogen, BMP2 directs gene transcription in recipient cells through suppressor of mothers against decapentaplegic-1/5/9 (SMAD1/5/9) transcription factors by binding to and signaling through complexes of type I and type II BMP receptors (*18*) **(Fig. 2D and fig. S3E)**. Primary pituitary cells responded to recombinant murine BMP2 (rmBMP2) with increased SMAD1/5 activation in a dose- and time-dependent manner **(Fig. 2E)**. Consistently, the canonical BMP target genes *Id1, Id2, Id3,* and *Smad6* were upregulated in rmBMP2-stimulated primary pituitary cells **(Fig. 2F)**.

We next treated the non-tumor-bearing wild-type mice with rmBMP2, testing whether BMP2 could recapitulate the same pattern of pituitary responses to CRC. Remarkably, intraperitoneal administration of rmBMP2 once daily for three consecutive days effectively activated pituitary endocrine function, as evidenced by increased levels of GH, prolactin, ACTH, β-endorphin, and LH **(Fig. 2H)**. To determine whether the *in vivo* effects of rmBMP2 on hormone secretion are mediated by a direct action on the pituitary, primary pituitary cells were stimulated with rmBMP2 *in vitro*. Consistent with the *in vivo* findings, rmBMP2 treatment led to a significant increase in hormone secretion **(fig. S3F)**. In addition, plasma levels of BMP2 were significantly elevated in both MC38 and AKPC tumor-bearing mouse models **(Fig. 2G)**. These results support BMP2 as a key factor mediating pituitary responses to CRC.

### Blockade of pituitary BMP signaling inhibits tumor progression

To investigate the functional relevance of the BMP2–pituitary axis in CRC development, we first performed pharmacological intervention in MC38 tumor-bearing mice **(Fig. 3, A and B)**. Remarkably, treatment with the extracellular BMP antagonist Noggin suppressed CRC-induced pituitary activation, as evidenced by reduced plasma levels of multiple pituitary hormones **(Fig. 3C and fig. S4A)**. Furthermore, recombinant murine Noggin (rmNoggin) treatment significantly reduced MC38 tumor burden **(Fig. 3, D and E)**. Importantly, neither rmBMP2 treatment nor pharmacological blockade of BMP signaling with rmNoggin affected MC38 cell viability *in vitro* **(fig. S4B)**.

**Fig. 3.**
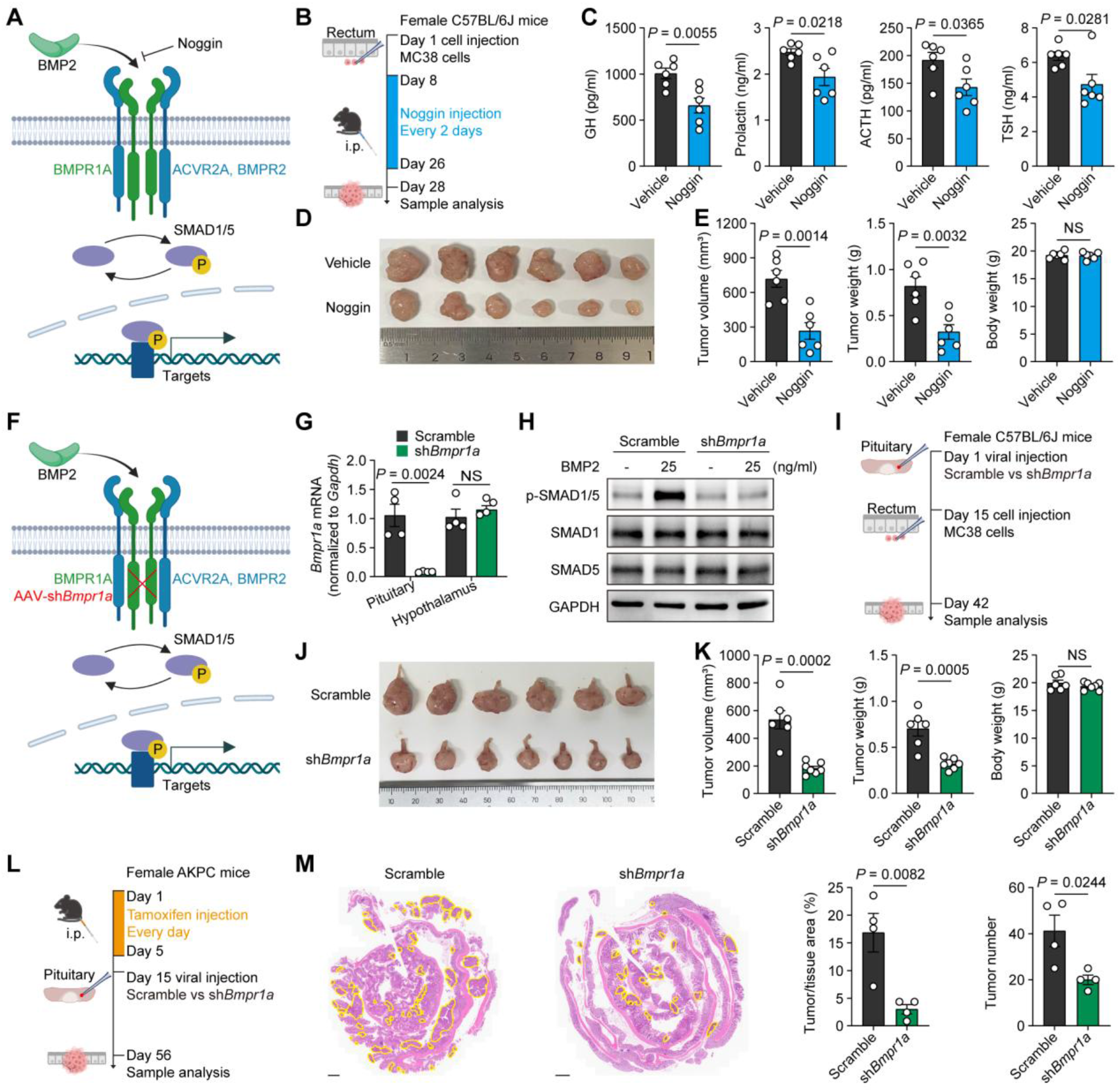
Blockade of pituitary BMP signaling inhibits tumor progression. (**A**) Illustration of pharmacological antagonism of BMP signaling by Noggin. (**B** to **E**) Pharmacological inhibition of the BMP signaling pathway in MC38 tumor-bearing mice. *n* = 6 mice per group. (B) Experimental design. (C) ELISA analysis of plasma pituitary hormones including GH, prolactin, ACTH and TSH in vehicle- and Noggin-treated mice. (D) Macroscopic morphology of rectal tumors. (E) Quantification of tumor volume, tumor weight, and body weight. (**F**) Illustration of pituitary-specific *Bmpr1a* knockdown via AAV-sh*Bmpr1a* to attenuate BMP receptor signaling. (**G**) qRT-PCR analysis to determine the *Bmpr1a* mRNA expression in the pituitary or hypothalamus of mice infected with AAV-sh*Bmpr1a* on day 28 after stereotaxic injection into the pituitary (*n* = 4). (**H**) Immunoblot analysis of cultured primary pituitary cells with control or *Bmpr1a* knockdown following treatment with 25 ng/mL rmBMP2 for 45 min. (**I** to **K**) Genetic knockdown of pituitary *Bmpr1a* in MC38 tumor-bearing mice. Scramble, *n* = 6 mice; sh*Bmpr1a*, *n* = 7 mice. (I) Experimental design. (J) Macroscopic morphology of rectal tumors. (K) Quantification of tumor volume, tumor weight, and body weight. (**L** to **M**) Pituitary-specific *Bmpr1a* knockdown in AKPC mice. *n* = 4 mice per group. (L) Experimental design. (M) Representative H&E staining of colorectal lesions from scramble- and sh*Bmpr1a*-treated AKPC mice (left). Yellow circles indicate tumor regions. Scale bars, 1 mm. Quantification of relative tumor tissue area (middle) and total tumor number (right). The data are presented as mean ± SEM. *P* values are determined by two-tailed unpaired Student’s *t* test (C, E, G, K and M). NS, not significant. Figure A, B, F, I and L created with BioRender.com.

In further support of the critical role of BMP2 signaling in the pituitary, we knocked down *Bmpr1a* in the pituitary using short hairpin RNA to disrupt receptor complex formation **(Fig. 3, F and G)**. Inhibition of *Bmpr1a* in the pituitary suppressed SMAD1/5 activation and reduced circulating hormone levels in MC38 tumor-bearing mice **(Fig. 3H and fig. S4C)**. Moreover, *Bmpr1a* knockdown led to a marked reduction in MC38 tumor growth, comparable to the effects of rmNoggin treatment **(Fig. 3, I to K)**.

We next examined the role of pituitary BMP signaling in an autochthonous CRC model. Two weeks after tumor initiation, adeno-associated virus–mediated knockdown of pituitary *Bmpr1a* was performed in AKPC mice to inhibit BMP signaling **(Fig. 3L)**. In this setting, pituitary *Bmpr1a* knockdown significantly suppressed CRC progression, as evidenced by reduced overall tumor burden and tumor number **(Fig. 3M)**. Together, these findings establish a crucial role for the BMP2–pituitary axis in CRC promotion.

### Pituitary promotes CRC progression via GHR^+^ CAFs

We next sought to identify potential peripheral targets of the CRC-induced pituitary endocrine response. To this end, we first examined the expression of pituitary hormone receptors in major immune and hematopoietic organs, including the bone marrow, spleen, and lymph nodes, using publicly available single-cell RNA-sequencing (scRNA-seq) datasets. *Mc5r* expression was enriched in a small population of common myeloid progenitors in the bone marrow, whereas *Prlr* was preferentially expressed by a relatively small population of splenic plasma cells. No comparably prominent receptor-expressing population was identified in the lymph nodes **(fig. S5)**. These findings suggested that pituitary hormones may engage discrete cell populations in distal immune and hematopoietic tissues.

We next asked whether pituitary hormones might also act directly within the TME. We therefore assessed the expression of pituitary hormone receptors in CRC by integrating and analyzing scRNA-seq data from MC38 tumor samples (*19–22*) **(Fig. 4A and fig. S6A)**. Cell types were identified based on the expression of classical cell type–specific marker genes **(fig. S6B)**. *Ghr* was the only hormone receptor abundantly expressed in the MC38 tumor, with high expression in cancer cells and fibroblasts but minimal expression in immune cell populations **(Fig. 4, B and C)**. Given the restricted distribution of candidate receptor-expressing populations in distal immune and hematopoietic tissues and the prominent expression of *Ghr* within the tumor, we focused subsequent functional studies on the GH–GHR axis in the TME.

**Fig. 4.**
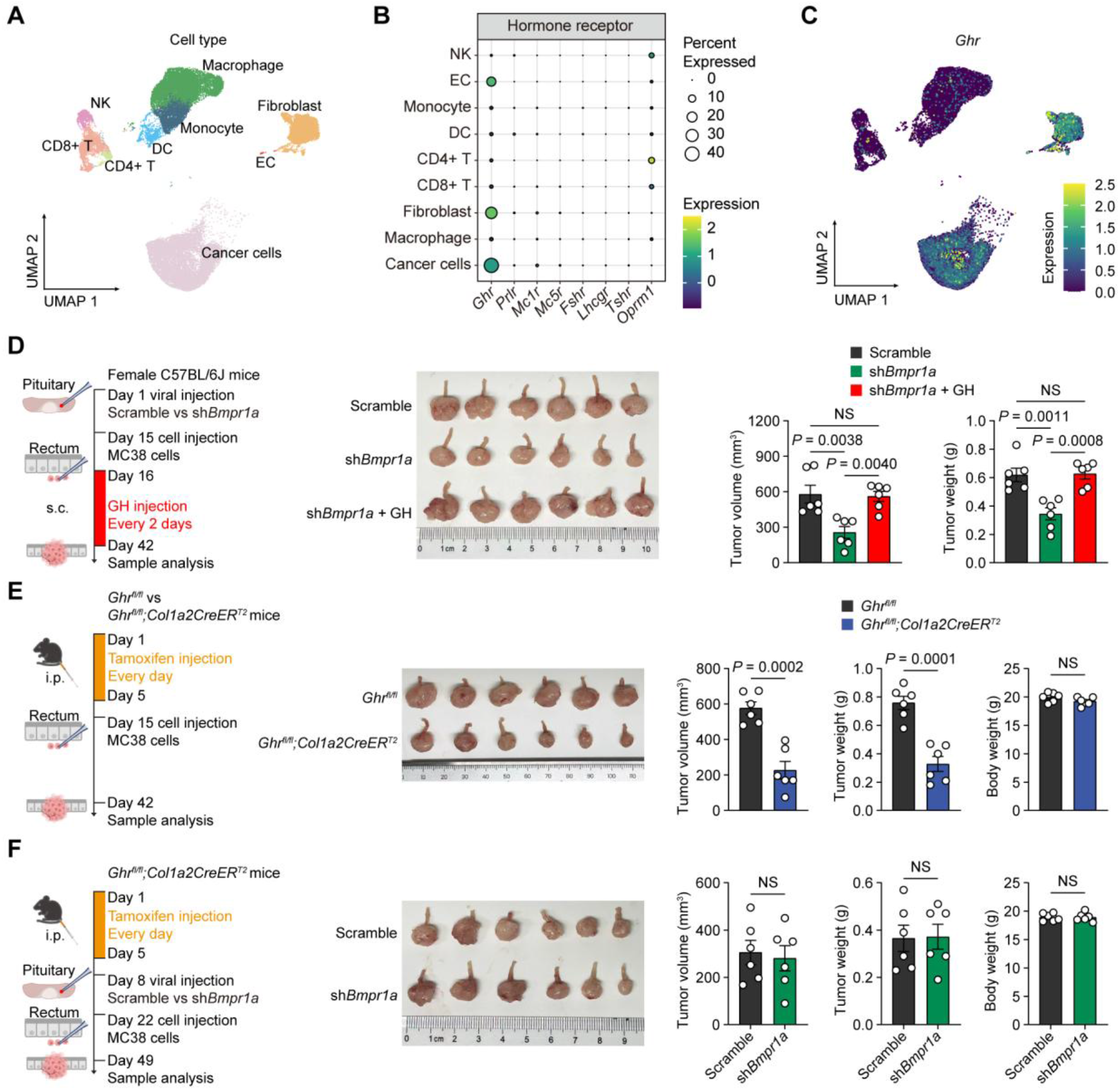
Pituitary promotes CRC progression via GHR^+^ CAFs. (**A** to **C**) scRNA-seq analysis of MC38 colorectal tumor tissues. (A) UMAP embedding showing major cell type clusters within the TME. (B) Dot plot showing the percentage of cells expressing and relative expression levels of indicated hormone receptors across all tumor cell subsets. *Mc2r*, *Mc3r*, and *Mc4r* were not detected. (C) Feature plot showing the cell-type-specific expression distribution of *Ghr* across tumor cell clusters. (**D**) Experimental setup (Created with BioRender.com.), representative rectal tumor images and tumor quantification in female C57BL/6J mice with pituitary scramble or sh*Bmpr1a* AAV injection, followed by MC38 orthotopic rectal tumor inoculation and subcutaneous rmGH supplementation every other day. Tumor burden was assessed by tumor volume and tumor weight (*n* = 6 per group). (**E**) Experimental setup (Created with BioRender.com.), representative rectal tumor images and tumor quantification in *Ghr^fl/fl^* and *Ghr^fl/fl^;Col1a2-CreER^T2^*mice orthotopically transplanted with MC38 tumor cells after tamoxifen induction. Tumor burden was assessed as tumor volume and tumor weight (*n* = 6 per group). (**F**) Experimental setup (Created with BioRender.com.), representative rectal tumor images and tumor quantification in *Ghr^fl/fl^;Col1a2-CreER^T2^*mice receiving pituitary scramble or sh*Bmpr1a* AAV prior to MC38 rectal tumor inoculation after tamoxifen induction. Tumor burden was assessed as tumor volume and tumor weight (*n* = 6 per group). The data are presented as mean ± SEM. *P* values are determined by two-tailed unpaired Student’s *t* test (E and F) and one-way ANOVA with Tukey’s multiple comparisons test (D). NS, not significant.

In line with the fibroblast-enriched expression pattern of *Ghr*, analysis of human CRC datasets revealed that *GHR* expression was highest in consensus molecular subtype 4 (CMS4) **(fig. S6C)**, the poor-prognosis, stroma-rich molecular subtype of CRC (*23*). Moreover, *Gh* expression was minimal in MC38 tumors **(fig. S6D)**. To examine the biological relevance of the GH–GHR axis in pituitary-mediated tumor promotion, we treated pituitary-specific *Bmpr1a* knockdown mice with recombinant murine GH (rmGH). Importantly, our results showed that activation of the GH–GHR axis via rmGH treatment significantly enhanced MC38 tumor growth in these mice, restoring tumor burden to a level comparable to that observed in mice with intact pituitary function **(Fig. 4D)**.

However, GH exerted minimal effects on MC38 cell growth *in vitro* (**fig. S6E**), suggesting that cancer cells are not direct cellular targets of this hormone. Given that fibroblasts also expressed *Ghr*, we hypothesized that GH promotes CRC progression through stromal fibroblasts. Fibroblasts are highly plastic and can transition from resting to activated states in response to microenvironmental cues, accompanied by transcriptional and functional changes in secretory, contractile, and extracellular matrix-remodeling programs (*24*). Accordingly, fibroblast-specific reclustering of 5,216 cells was followed by stratification according to their activation state, yielding resting and activated fibroblast populations **(fig. S6F)**. Both populations retained fibroblast identity, as indicated by *Dcn* expression, whereas activated fibroblasts exhibited a coordinated increase in activation-associated markers, including *Acta2*, *Pdgfra*, *Fap*, *Lrrc15* and *Pdpn* (*25*) **(fig. S6G)**. Notably, *Ghr* expression was preferentially enriched in activated fibroblasts **(fig. S6G)**, supporting activated fibroblasts as a potential stromal target of GH signaling. We further evaluated the relationship between *GHR* expression and stromal/CAF infiltration in CRC. Across TCGA COAD and READ cohorts, *GHR* expression positively correlated with stromal scores and CAF abundance estimated by EPIC, MCP-counter, xCell, and TIDE **(fig. S6H)**. Consistently, *GHR* expression was associated with multiple fibroblast activation markers, including *ACTA2*, *FAP*, *PDGFA*, *PDPN*, and *LRRC15* **(fig. S6I)**. We next isolated primary CAFs from MC38 tumors (CAF-1) and AKPC tumors (CAF-2), which were cultured as primary cells and validated by western blotting and immunofluorescence (IF) **(fig. S7, A and B)**. GHR expression in these primary CAF populations was verified at the protein level **(fig. S7C)**.

To test whether GH promotes CRC progression via GHR expressed in CAFs *in vivo*, we established MC38 tumors in mice with fibroblast-specific *Ghr* deletion (*Ghr^fl/fl^;Col1a2-CreER^T2^*mice) **(Fig. 4E and fig. S7, D and E)**. *Col1a2-CreER^T2^* has been previously validated for efficient and selective fibroblast recombination with minimal off-target effects (*26*), and our integrated mouse MC38 tumor scRNA-seq dataset further confirmed preferential *Col1a2* expression in fibroblasts **(fig. S7F)**. We further confirmed that *Ghr* expression was significantly reduced *ex vivo* in tamoxifen-treated *Ghr^fl/fl^;Col1a2-CreER^T2^*CAFs **(fig. S7G)**. Importantly, mice with fibroblast-specific *Ghr* deletion exhibited markedly reduced tumor burden compared with *Ghr^fl/fl^* littermate controls when both were orthotopically transplanted with wild-type tumor cells **(Fig. 4E)**, indicating that the GH–GHR axis acts in a CAF-dependent manner to control CRC growth. In addition to reduced tumor growth, fibroblast-specific *Ghr* ablation significantly prolonged median survival, attenuated tumor-associated weight loss, and preserved nesting scores, indicating improved activity and overall welfare **(fig. S7, H to J)**. Consistent with the clinical relevance of these findings, higher *GHR* expression was associated with poorer overall survival in patients with CMS4 colon cancer, a subtype associated with poor therapeutic response and adverse prognosis **(fig. S7K)**. Together, these results reveal that GHR^+^ CAFs play a key role in CRC progression. Furthermore, *Ghr* deficiency in fibroblasts abolished the tumor-inhibitory effect of pituitary *Bmpr1a* depletion **(Fig. 4F)**, indicating that the GH–GHR axis mediates the downstream effects of the BMP2–pituitary axis in CRC.

### GH–GHR signaling induces a senescence-like phenotype in CAFs

To investigate how CAFs contribute to CRC progression, we performed RNA-seq analysis on CAF-1 cells treated with or without GH. GH stimulation induced distinct gene expression patterns compared with vehicle control **(Fig. 5, A and B)**. Enriched signaling pathway analysis revealed increased senescence-associated features in GH-treated CAF-1 cells **(Fig. 5C)**, accompanied by elevated expression of mRNAs encoding multiple senescence-associated secretory phenotype (SASP) factors—including interleukins, cytokines, chemokines, and proteases—as previously summarized (*27*) **(Fig. 5D)**.

**Fig. 5.**
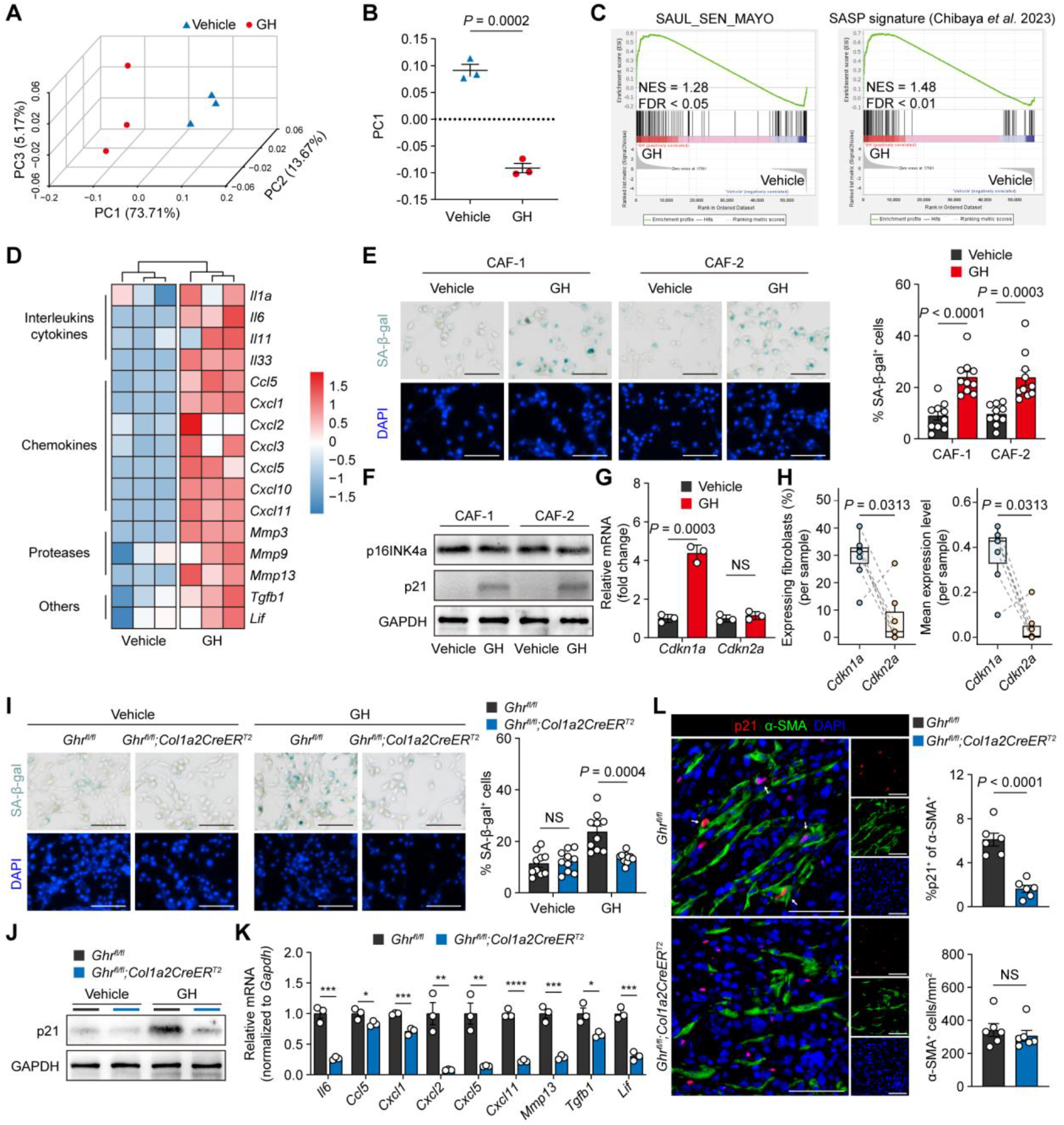
GH–GHR signaling induces a senescence-like phenotype in CAFs. (**A** to **D** and **G**) Bulk RNA-seq analysis of CAF-1 cells treated with vehicle or GH (500 ng/mL) for 7 d. (A) Three-dimensional PCA plot. (B) Quantification of PC1 scores from the PCA analysis in (A). (C) Gene set enrichment analysis (GSEA) for senescence and SASP signatures. NES and false discovery rate (FDR) values are labeled for each gene set. (D) Heatmap showing relative expression of representative SASP-related genes. (**E**) Representative images and quantitative analysis of the percentage of SA-β-gal-positive cells in CAF-1 and CAF-2 treated with vehicle or 500 ng/mL GH for 7 d (*n* = 10 independent cultures). Scale bar, 100 μm. (**F**) Immunoblot analysis of p16INK4a and p21 in CAF-1 and CAF-2 treated with vehicle or 500 ng/mL GH for 7 d, with GAPDH as loading control. (G) Relative mRNA expression of *Cdkn1a* and *Cdkn2a* based on FPKM values. (**H**) Paired comparisons of *Cdkn1a* and *Cdkn2a* expression in the fibroblast population from the integrated MC38 tumor scRNA-seq datasets presented in Figure 4A. Left, percentages of fibroblasts with detectable *Cdkn1a* or *Cdkn2a* expression in each tumor sample. Right, mean normalized expression levels of *Cdkn1a* and *Cdkn2a* in fibroblasts from each tumor sample. Each pair of connected points represents one independent tumor sample (*n* = 7). Box plots show the median and interquartile range. *P* values were determined using two-tailed paired Wilcoxon signed-rank tests. (**I**) Representative images and quantitative analysis of the percentage of SA-β-gal-positive cells in primary CAFs isolated from *Ghr^fl/fl^* and *Ghr^fl/fl^;Col1a2-CreER^T2^* mice after treatment with vehicle or 500 ng/mL GH for 7 d (*n* = 10 independent cultures). Scale bar, 100 μm. (**J**) Immunoblot analysis of p21 in primary CAFs isolated from *Ghr^fl/fl^* and *Ghr^fl/fl^;Col1a2-CreER^T2^* mice after treatment with vehicle or 500 ng/mL GH for 7 d, with GAPDH as loading control. (**K**) qRT-PCR analysis detecting relative mRNA expression of representative SASP-related genes in primary CAFs isolated from *Ghr^fl/fl^*and *Ghr^fl/fl^;Col1a2-CreER^T2^* mice following GH treatment for 7 d (*n* = 3). (**L**) Representative co-IF images and quantification of MC38 tumors from *Ghr^fl/fl^*and *Ghr^fl/fl^;Col1a2-CreER^T2^* mice (*n* = 6). Scale bars, 50 μm. White arrows indicate p21^+^ α-SMA^+^ CAFs. The upper panel shows the percentage of p21^+^ cells among α-SMA^+^ CAFs, and the lower panel displays the density of total α-SMA^+^ CAFs per mm^2^. The data are presented as mean ± SEM. *P* values are determined by two-tailed unpaired Student’s *t* test (B, E, G, I, K and L).

To validate this observation, the expression of senescence markers was examined in CAFs after GH treatment. Quantification of senescence-associated β-galactosidase (SA-β-Gal) staining demonstrated a significant increase in SA-β-Gal positivity in GH-treated CAFs compared with untreated controls **(Fig. 5E)**. Immunoblotting confirmed increased p21 protein expression without a significant change in p16INK4a, consistent with the RNA-seq results from GH-treated CAF-1 cells **(Fig. 5, F and G)**. Additionally, analysis of the fibroblast population in the integrated MC38 tumor scRNA-seq dataset showed that *Cdkn1a* was detected in a greater proportion of cells and exhibited higher mean expression than *Cdkn2a* across independent tumor samples **(Fig. 5H)**, supporting the predominance of a *Cdkn1a*-associated transcriptional program in MC38 tumor fibroblasts.

Notably, GH-induced senescence phenotypes were abolished in *Ghr*-deficient CAFs. Both SA-β-Gal activity and p21 accumulation were reduced in the absence of GHR **(Fig. 5, I and)**. Similarly, the upregulation of SASP factors was also dependent on GHR **(Fig. 5K)**. In agreement with these findings, fibroblast-specific *Ghr* deletion in MC38 orthotopic tumors led to a significant reduction in nuclear p21 expression in α-SMA^+^ CAFs **(Fig. 5L)**, indicating that a similar senescence program was also induced *in vivo*. Together, these findings support a GHR-dependent senescence-like response to GH in CAFs.

We then investigated the mechanism by which the GH–GHR axis induces a senescence-like phenotype in CAFs. GHR belongs to the class I cytokine receptor family and signals, at least in part, through the Janus-family tyrosine kinase 2 (JAK2)–signal transducer and activator of transcription 5 (STAT5) pathway (*28*). Previous studies have implicated STAT5 signaling in the regulation of cell-cycle inhibitors, including p21, and in cellular senescence (*29, 30*). We therefore examined whether JAK2–STAT5 signaling mediates the senescence-like response to GH in CAFs. CAF-1 cells were first treated with fedratinib, a JAK2 inhibitor (*31*), which dose-dependently reduced GH-induced p21 protein expression **(fig. S8A)**. Similarly, treatment with STAT5-IN-1, a potent and selective STAT5 inhibitor (*32*), resulted in a dose-dependent suppression of GH-induced p21 protein expression **(fig. S8B)**. Consistent with these protein-level findings, treatment with fedratinib or STAT5-IN-1 also decreased GH-induced *Cdkn1a* transcript levels **(fig. S8C)**. Moreover, inhibition of either JAK2 or STAT5 attenuated GH-induced SA-β-gal activity in CAF-1 cells, suggesting that activation of the JAK2–STAT5–p21 axis contributes to the GH-induced senescence-like phenotype in CAFs **(fig. S8D)**.

SOCS1 plays a critical role in cytokine signaling and links STAT5 activation to cellular senescence by promoting p53 activation (*30*). We found that GH treatment significantly increased *Socs1* expression and enriched the p53 signaling pathway in CAF-1 cells **(fig. S8, E and F)**. Together, these findings suggest that GH–GHR signaling promotes a senescence-like phenotype in CAFs, at least in part, through activation of the JAK2–STAT5–p21 axis.

### p21^+^ CAFs are required for GH–GHR axis-mediated tumor promotion and predict poor prognosis

To investigate the *in vivo* functions of senescent CAFs in CRC, we induced fibroblast-specific knockdown of *Cdkn1a* or *Cdkn2a* by rectal submucosal injection of AAV-DIO-sh*Cdkn1a* or AAV-DIO-sh*Cdkn2a* into female *Col1a2-CreER^T2^* mice, followed by orthotopic CRC tumor implantation **(Fig. 6A and fig. S9, A and B)**. To verify efficient *Cdkn1a* knockdown in fibroblasts *in vivo*, we performed co-immunofluorescence staining for p21 together with α-SMA, a well-established marker of activated CAFs in CRC (*33*). Conditional knockdown (cKD) of *Cdkn1a* markedly reduced p21 expression in α-SMA^+^ CAFs **(Fig. 6B)**. Functionally, *Cdkn1a* cKD suppressed MC38 tumor growth and prolonged overall survival, supporting a tumor-promoting role for p21^+^ senescent CAFs in this model **(Fig. 6, C and D)**. Consistently, analysis of the TCGA COAD and READ datasets showed that high *CDKN1A* expression was associated with poor prognosis in patients with CRC (**fig. S9C**). Notably, in the absence of p21^+^ CAFs, CAF-specific *Ghr* ablation no longer conferred additional tumor suppression or survival benefit **(Fig. 6, C and D)**, indicating that p21 is required for the GHR^+^ CAF-mediated CRC progression. By contrast, cKD of *Cdkn2a* was insufficient to abrogate the tumor-inhibitory effect induced by CAF-specific *Ghr* depletion **(fig. S9B)**. Together, these results reveal that p21, but not p16INK4a, is required for GH–GHR axis-mediated regulation of CAFs and CRC progression.

**Fig. 6.**
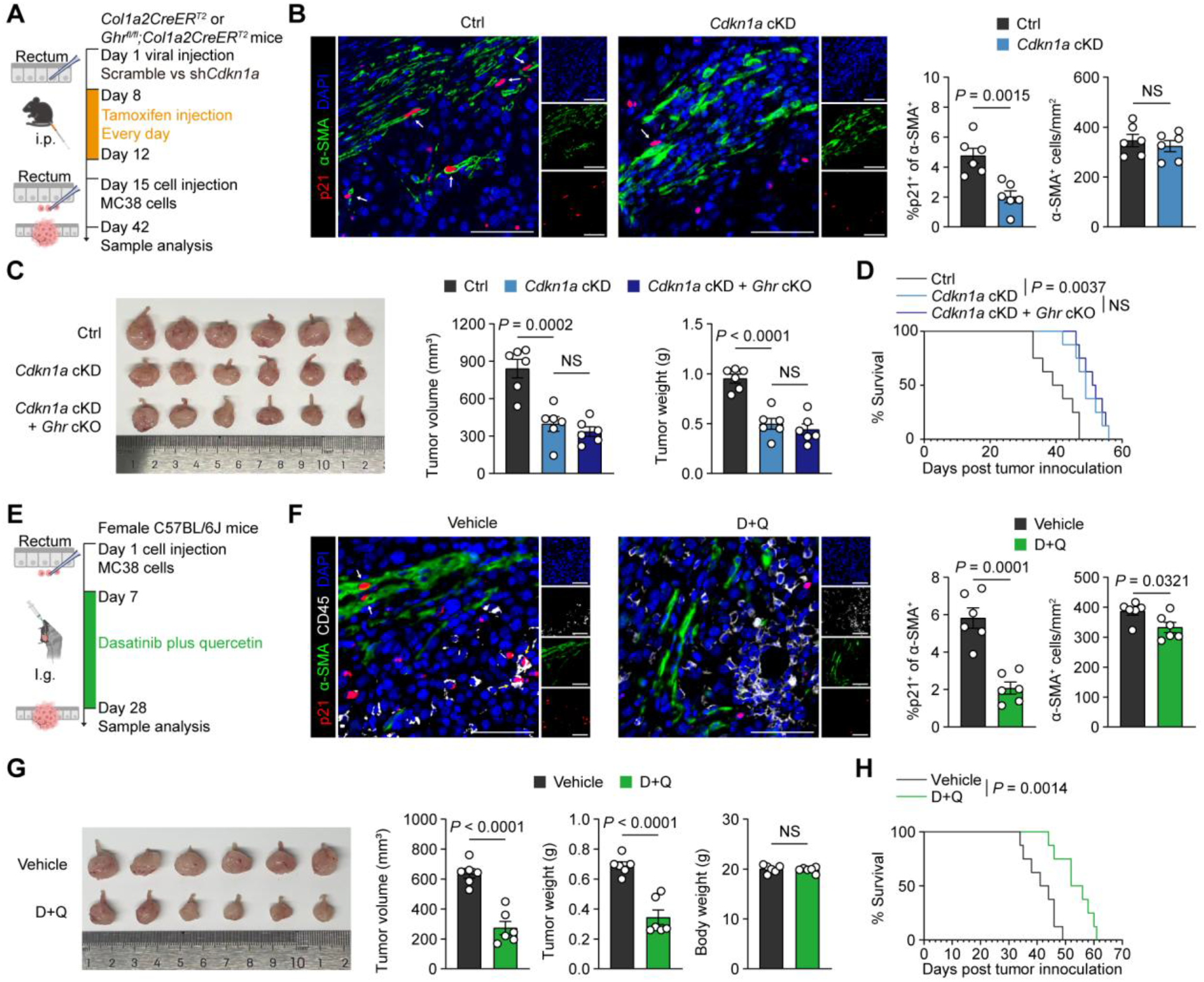
p21⁺ CAFs are required for GH–GHR axis-mediated tumor promotion. (**A** to **D**) Cre-dependent knockdown of *Cdkn1a* in rectal fibroblasts suppresses MC38 tumor growth, and this effect is not further enhanced by concomitant *Ghr* deletion. *n* = 6 mice per group. (A) Experimental design. *Col1a2-CreER^T2^* or *Ghr^fl/fl^;Col1a2-CreER^T2^* mice received rectal viral injection of AAV9-DIO-miR30-shNC (Ctrl) or AAV9-DIO-miR30-sh*Cdkn1a* (*Cdkn1a* cKD), followed by tamoxifen administration and orthotopic MC38 tumor inoculation. (B) Representative co-IF images and quantification of MC38 tumors from control and *Cdkn1a* cKD mice. Scale bars, 50 μm. White arrows indicate p21^+^ α-SMA^+^ CAFs. The left graph shows the percentage of p21^+^ cells among α-SMA^+^ CAFs, and the right graph shows the density of total α-SMA^+^ CAFs per mm^2^. (C) Representative rectal tumor images and quantification of tumor volume and tumor weight in control, *Cdkn1a* cKD, and *Cdkn1a* cKD + *Ghr* cKO mice. (D) Kaplan–Meier survival plots of control, *Cdkn1a* cKD, and *Cdkn1a* cKD + *Ghr* cKO mice following orthotopic inoculation of MC38 cells (log-rank test). (**E** to **H**) Pharmacological clearance of senescent cells suppresses rectal tumor growth and prolongs survival in MC38 tumor-bearing mice. *n* = 6 mice per group. (E) Experimental design. Female C57BL/6J mice were orthotopically inoculated with MC38 cells and treated with vehicle or dasatinib plus quercetin (D+Q) from day 7 to day 27, followed by sample collection on day 28. (F) Representative co-IF images and quantification of MC38 tumors from vehicle- and D+Q-treated mice. Scale bars, 50 μm. White arrows indicate p21^+^ α-SMA^+^ CAFs, yellow arrows indicate p21^+^ CD45^+^ immune cells. The left graph shows the percentage of p21^+^ cells among α-SMA^+^ CAFs, and the right graph shows the density of total α-SMA^+^ CAFs per mm^2^. (G) Representative rectal tumor images and quantification of tumor volume, tumor weight and body weight in vehicle- and D+Q-treated mice. (H) Kaplan–Meier survival plots of MC38 tumor-bearing mice treated with vehicle or D+Q (log-rank test). The data are presented as mean ± SEM. *P* values are determined by two-tailed unpaired Student’s *t* test (B, F and G) and one-way ANOVA with Tukey’s multiple comparisons test (C). Figures A and E were created with BioRender.com.

As complementary pharmacological validation, we treated tumor-bearing wild-type mice with dasatinib plus quercetin (D+Q), a canonical senolytic regimen (*34*). D+Q treatment significantly decreased the abundance of p21^+^α-SMA^+^ senescent CAFs, reduced tumor burden, and extended overall survival **(Fig. 6, E to H)**. However, D+Q treatment also reduced p21^+^CD45^+^ cells **(fig. S9D)**, suggesting that its antitumor effect was not exclusively attributable to senescent CAF depletion. This reduction may result from several contributors, including direct elimination of senescent immune cells by D+Q and secondary remodeling of immune-cell infiltration following senescent CAF depletion. Moreover, D+Q directly impaired MC38 cell viability *in vitro* **(fig. S9E)**. Therefore, given the broad senolytic activity and tumor cell–intrinsic effects of D+Q, we could not definitively attribute the reduced tumor burden solely to senescent CAF clearance and did not use this pharmacological approach for further causal verification.

### Human pituitary metabolic activity is elevated in patients with CRC and correlates with stromal p21 expression and tumor burden

Lastly, we investigated the clinical relevance of the pituitary–stromal axis in patients with CRC. Given that pituitary ^18^F-fluorodeoxyglucose (^18^F-FDG) uptake on positron emission tomography/computed tomography (PET/CT) has been associated with endocrine activity (*35*), we analyzed ^18^F-FDG PET/CT data from 50 non-malignant controls and 52 patients with CRC **(table S4)**. To improve comparability across individuals, we calculated the pituitary standardized uptake value ratio (SUVR) as an indicator of pituitary glucose metabolic activity (*36*). Pituitary SUVR was significantly higher in patients with CRC than in non-malignant controls **(Fig. 7A)**.

**Fig. 7.**
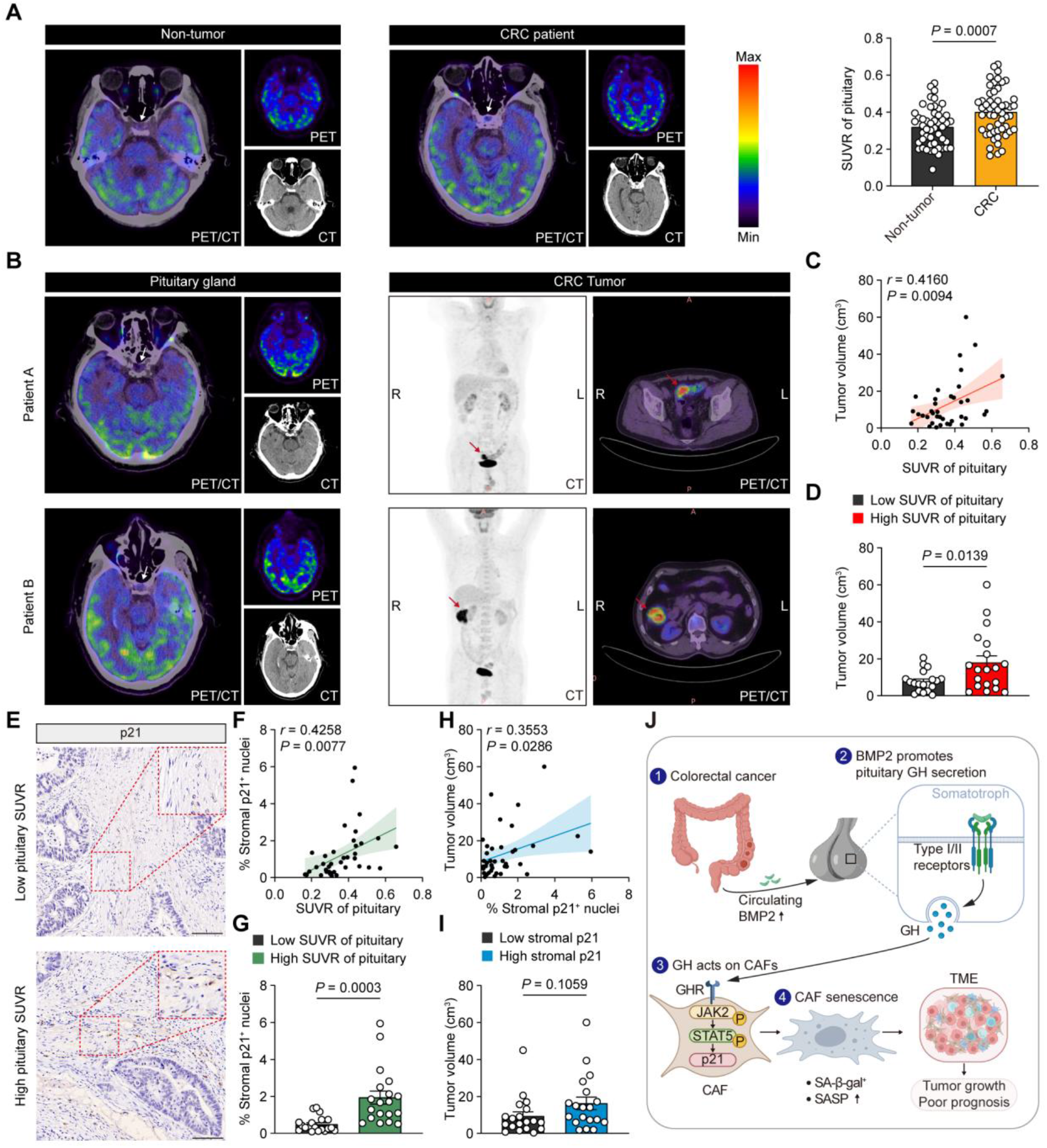
Human pituitary metabolic activity is elevated in patients with CRC and correlates with stromal p21 expression and tumor burden. **(A)** Representative ^18^F-FDG PET/CT images of the pituitary gland in non-malignant controls (*n* = 50) and patients with CRC (*n* = 52), and comparison of pituitary ^18^F-FDG SUVR between the two groups. White arrows indicate the pituitary gland. The color bar indicates ^18^F-FDG standardized uptake value. (B) Representative ^18^F-FDG PET/CT images of the pituitary gland and corresponding primary colorectal tumors from patients with CRC. White arrows indicate the pituitary gland, and red arrows indicate the primary tumor lesions. (C) Correlation analysis between pituitary ^18^F-FDG SUVR and tumor volume. The solid line indicates the linear regression fit, and the shaded area represents the 95% confidence interval (*n* = 38). Pearson correlation analysis. (D) Comparison of tumor volume between the low (*n* = 19) and high (*n* = 19) pituitary ^18^F-FDG SUVR groups. (E) Representative IHC staining of p21 in CRC tissues from patients with low or high pituitary SUVR. Arrows indicate stromal cells with nuclear p21 expression. Scale bars, 100 μm. (**F**) Correlation analysis between pituitary ^18^F-FDG SUVR and the percentage of stromal cells with nuclear p21 expression. The solid line indicates the linear regression fit, and the shaded area represents the 95% confidence interval (*n* = 38). Pearson correlation analysis. (**G**) Comparison of the percentage of stromal cells with nuclear p21 expression between the low (*n* = 19) and high (*n* = 19) pituitary ^18^F-FDG SUVR groups. (**H**) Correlation analysis between the percentage of stromal cells with nuclear p21 expression and tumor volume. The solid line indicates the linear regression fit, and the shaded area represents the 95% confidence interval (*n* = 38). Pearson correlation analysis. (**I**) Comparison of tumor volume between the low (*n* = 19) and high (*n* = 19) stromal p21 groups. (**J**) CRC elevates circulating BMP2, thereby enhancing pituitary GH secretion. Elevated GH activates GHR–JAK2–STAT5 signaling in CAFs, leading to increased p21 expression and CAF senescence-like state. This senescent state is associated with increased SA-β-gal activity and SASP, promoting tumor growth and poor prognosis. The data are presented as mean ± SEM. *P* values are determined by two-tailed unpaired Student’s *t* test (A, D, G and I). Figure J created with BioRender.com.

Among the 52 patients with CRC, 38 underwent surgical resection following their initial diagnosis, allowing pathological assessment of tumor volume and collection of matched tumor specimens **(table S4)**. Within this surgically resected subset, pituitary SUVR was positively correlated with tumor volume **(Fig. 7, B to D)**. By contrast, pituitary SUVR was not associated with tumor SUVmax **(fig. S10, A and B)**, suggesting that increased pituitary metabolic activity is more closely related to overall tumor burden than to the intrinsic glucose metabolic intensity of the tumor.

Analysis of the corresponding tumor specimens further revealed a stromal association. Immunohistochemical (IHC) analysis showed that the proportion of stromal cells with nuclear p21 expression was positively correlated with both pituitary SUVR and tumor volume **(Fig. 7, E to I)**. In contrast, nuclear p21 expression in tumor cells was not significantly associated with either pituitary SUVR or tumor volume **(fig. S10, C to F)**. Thus, the association between p21 expression, pituitary metabolic activity, and tumor burden was preferentially evident in the stromal compartment rather than in tumor cells. Collectively, these findings extend the pituitary–stromal axis identified in our experimental models to human CRC and support its clinical relevance.

## Discussion

Tumors can perturb systemic endocrine homeostasis, and accumulating evidence suggests that such neuroendocrine alterations can contribute to disease progression (*4, 8*). However, how peripheral tumors engage the pituitary and whether this response functionally remodels the local TME remain poorly understood. Here, we show that CRC activates a tumor–pituitary–stromal axis in which elevated circulating BMP2 enhances pituitary endocrine output, and pituitary-derived GH acts on GHR-expressing CAFs to induce a p21-dependent senescence-like, tumor-promoting state **(Fig. 7J)**. Disruption of pituitary BMP signaling, fibroblast-specific *Ghr* deletion, or fibroblast-specific *Cdkn1a* knockdown markedly restrained tumor progression, supporting a functional role for this endocrine–stromal pathway in CRC progression.

The present study identifies circulating BMP2 as an upstream mediator linking CRC to pituitary endocrine activation. At least two clinical studies have reported elevated circulating BMP2 in patients with cancer, with higher serum levels associated with disease progression and poor outcome in gastric cancer and advanced non-small-cell lung cancer (*37, 38*). In the colonic epithelium, BMP2 has been reported to exert tumor-suppressive effects by promoting differentiation and apoptosis while restraining proliferation (*39*). However, this function is highly context dependent. Loss of SMAD4 can switch BMP signaling from tumor suppressive to metastasis promoting (*40*), and fibroblast-derived BMP2 has been shown to enhance the invasion and metastasis of SMAD4-deficient CRC cells (*41*). In the MC38 model, BMP2 modulation had no detectable effect on tumor cell viability in vitro. Rather, BMP2 promoted CRC progression through a host endocrine response involving the pituitary. BMP signaling plays important roles in pituitary development and endocrine cell specification (*42*). Our data suggest that BMP-responsive signaling remains operative in the adult pituitary and can be engaged by CRC-associated circulating BMP2 to enhance endocrine activity. The cellular source of the increased circulating BMP2, as well as the mechanisms responsible for its elevation during CRC, remain to be determined.

Cross-tissue receptor analysis suggests that CRC-induced pituitary remodeling may engage multiple peripheral compartments. Candidate pituitary hormone receptors showed relatively restricted distributions in distal immune and hematopoietic tissues, whereas *Ghr* was prominently expressed within the MC38 TME, particularly in cancer cells and fibroblasts. This intratumoral enrichment of *Ghr* prompted further mechanistic investigation of the GH–GHR axis in the TME. Endocrine actions in distal organs or parallel effects mediated by other pituitary hormones remain possible.

The findings indicate that pituitary-derived endocrine GH contributes to the stromal response in CRC. Non-pituitary GH has been detected in multiple peripheral tissues and can exert potent autocrine or paracrine effects (*43*), but its production is generally low and does not substantially contribute to circulating GH levels (*44*). Thus, the increase in circulating GH observed in mouse models of CRC is most consistent with enhanced pituitary endocrine output. This does not exclude a contribution from locally produced GH within the TME. Local GH can increase with aging and DNA damage, and has been detected in both colonic epithelial cells and fibroblasts under such conditions (*45*). Although single-cell analysis did not detect appreciable *Gh* expression across cell populations in MC38 tumors, this observation alone cannot exclude low-level, transient, or context-dependent autocrine/paracrine GH production. Fibroblast-specific *Ghr* deletion shows that GH sensing by CAFs is required, while pituitary perturbation and GH rescue support a functional contribution of pituitary-derived GH to this stromal response. Pituitary-derived GH can stimulate hepatic IGF-1 production (*46*), and IGF-1 may also be produced locally within peripheral tissues and the TME (*47, 48*). Our findings support a direct action of GH through GHR in CAFs, while additional effects of systemic or local IGF-1 cannot be excluded.

Here, we identify GH as an upstream endocrine signal that promotes a p21-dependent senescence-like state in CRC CAFs and thereby promotes tumor growth. Senescence is highly heterogeneous, with p21 and p16INK4a variably engaged across cell lineages, inducing stimuli, and tissue contexts (*49*). Previous studies have identified tumor-promoting p16INK4a^+^ senescent populations within myeloid and fibroblast lineages in several cancer models (*50–53*), whereas more recent work in prostate cancer revealed a distinct p21^+^ stromal senescent population characterized by a prominent inflammatory SASP and tumor-promoting activity (*54*). Although p21 can precede p16INK4a induction in some senescence settings (*55, 56*), single-cell studies have also shown that p21^+^ and p16INK4a^+^ cells can occupy distinct, minimally overlapping states with different trajectories and secretory profiles (*57*).

While our data place the GHR–JAK2–STAT5 axis upstream of p21 induction, how GH signaling induces and maintains the broader senescent phenotype remains unresolved. Senescence involves coordinated remodeling of cell-cycle control, chromatin state, metabolism, and intercellular signaling (*58*), and p21 itself has been shown to regulate secretory remodeling through Rb-dependent transcriptional programs (*59*). Thus, p21 may contribute not only to growth arrest but also to the functional properties of GH-induced senescent CAFs. The SASP represents a major determinant of the context-dependent effects of senescence in cancer, and its composition can vary with the duration and nature of the inducing stimulus, cell identity, and tissue environment (*49, 58*). Although our data show that GH-responsive CAFs acquire a senescence-associated secretory program, the specific SASP components that contribute to tumor progression, how they evolve over time, and how they reshape immune and tumor-cell behavior in CRC remain to be defined.

The enrichment of GHR in CMS4 CRC may have particular translational relevance. CMS4 tumors are characterized by prominent stromal activation and poor clinical outcome, and our analysis showed increased *GHR* expression in this subtype, with higher *GHR* expression associated with worse survival. Importantly, pharmacological inhibition of GHR is already feasible. The FDA-approved GHR antagonist pegvisomant has shown antitumor activity in several preclinical cancer models, including breast, prostate, gastric, and hepatocellular cancers, where it suppresses tumor growth and, in some settings, enhances therapeutic response (*60–64*). Given our findings, patients with GHR-high or CMS4-type CRC may therefore represent a rational population in which GHR-directed therapy, alone or in combination with standard treatment, could be evaluated in future studies.

In summary, our findings identify the pituitary as a functional intermediary between CRC-associated systemic signals and peripheral effector compartments. Pituitary-derived GH acts on GHR-positive CAFs to promote a p21-dependent, tumor-promoting senescence-like state, revealing a mechanism by which a distant endocrine organ can shape tumor progression. This work expands the concept of tumor–host communication beyond local interactions and suggests that targeting systemic endocrine–stromal signaling may offer a strategy to disrupt host physiological programs exploited by CRC.

## Materials and Methods

### Mice

C57BL/6J mice were purchased from GemPharmatech Co., Ltd. (Shanghai, China). *CDX2P-CreER^T2^* (022390) and *Apc^15lox^* (029275) mouse strains were kindly provided by Prof. Yujun Hao from the Shanghai Cancer Institute, Renji Hospital, Shanghai Jiao Tong University School of Medicine. Conditional *Kras^LSL-G12D^* and *Trp53^R172H^* mice were obtained from The Jackson Laboratory, as described previously (*65*). *Ghr^flox^* and *Col1a2-CreER^T2^*mice were generated by GemPharmatech Co., Ltd. using CRISPR/Cas9-mediated genome editing. Mice were maintained under specific pathogen-free conditions at 24 ± 1°C and 50–70% relative humidity on a 12-h light/dark cycle, with lights on from 08:00 to 20:00. Standard chow and water were provided ad libitum, and mice were group-housed at no more than five animals per cage. Animals were monitored for body weight, tumor burden, and general health. The maximum permitted tumor burden was 2,000 mm^3^ or a maximal tumor diameter of 2 cm, and mice were euthanized earlier if signs of distress or deterioration in general condition were observed. All animal procedures were conducted in accordance with the Regulations for the Administration of Affairs Concerning Experimental Animals in China and were approved by the Ethics Committee of Ren Ji Hospital, affiliated with Shanghai Jiao Tong University (No. RA-2021-387).

### Cell lines and cell culture

The murine colon cancer cell line MC38-luc was obtained from Quicell Biotech (Shanghai, China). MC38 cells were cultured in Dulbecco’s modified Eagle’s medium (DMEM; 11965092, Gibco, USA) supplemented with 10% fetal bovine serum (FBS; A5670701, Gibco) and 1% penicillin–streptomycin (15140148, Gibco) at 37°C in a humidified atmosphere containing 5% CO_2_. Cell lines were authenticated by short tandem repeat profiling and tested for mycoplasma contamination upon receipt using a Mycoplasma PCR Detection Kit (BL1469A, Biosharp).

### Human samples

The use of pathological specimens and the review of relevant patient records were approved by the institutional research ethics committee and conducted in accordance with the Declaration of Helsinki and its subsequent amendments or comparable ethical standards. A retrospective cohort comprising 52 patients with CRC and 50 non-malignant controls who underwent clinical ^18^F-FDG PET/CT examination was analyzed. PET/CT images were obtained from Shanghai Tenth People’s Hospital. Among the 52 patients with CRC, tumor specimens were available from 38 patients who subsequently underwent surgical resection and were used for stromal p21 analysis. All clinical specimens were collected with approval from the Ethics Review Committee of Ren Ji Hospital (No. KY2021-120-B). Written informed consent was obtained from all patients.

### Orthotopic MC38 tumor model

Female C57BL/6J mice aged 6–8 weeks were used for the standard orthotopic MC38 tumor model. For experiments involving inducible genetically engineered mice, tumor cell transplantation was performed at the indicated time points after tamoxifen-induced recombination. MC38 cells were harvested, washed with PBS, and resuspended at 2 × 10^5^ cells in 20 μL PBS per mouse. Mice were immobilized in well-ventilated Perspex tubes, and tumor cells were injected into the submucosa of the posterior rectal wall, approximately 0.5 cm from the anal verge, using a Hamilton microsyringe under stereomicroscopic guidance. Successful injection was indicated by the formation of a transparent submucosal bulge. Control mice received an equal volume of PBS. For bioluminescence imaging, mice bearing luciferase-expressing MC38 tumors were anesthetized with isoflurane and administered D-luciferin sodium salt intraperitoneally (150 mg/kg; SB-D1007, ShareBio, Shanghai, China). Ten minutes later, bioluminescence signals were acquired using an IVIS Spectrum imaging system (PerkinElmer) and quantified using Living Image software. At the indicated endpoints, tumors were excised and weighed. Tumor volume was calculated as longest diameter × vertical diameter × height / 2 (mm^3^).

### Brain-region dissection and bulk RNA sequencing

At endpoint, brains were rapidly removed and dissected into the cerebrum, cerebellum, hindbrain, hypothalamus, and thalamus with midbrain, together with the pituitary gland. Total RNA was extracted from each tissue and quality assessed before library construction. Poly(A)-enriched mRNA libraries were subjected to paired-end sequencing on a NovaSeq 6000 sequencing platform (Illumina). Raw reads were processed using fastp and aligned to the mouse reference genome GRCm39 (Ensembl release 108) using HISAT2. Gene expression was quantified using StringTie, normalized using the trimmed mean of M values (TMM) method, and represented as fragments per kilobase of transcript per million mapped reads (FPKM). PCA was performed in R on Hellinger-transformed expression data using decostand in the vegan package and prcomp, and visualized using scatterplot3d (v0.3-44). Differential gene-expression analysis was performed using edgeR, with differentially expressed genes defined as P < 0.05 and fold change ≥ 2 or ≤ 0.5. Sequencing data have been deposited in the NCBI BioProject database under accession number PRJNA1512587.

### Genetically engineered mice and genotyping

To generate AKPC mice, *CDX2P-CreER^T2^* mice were crossed with *Apc^15lox^* mice to obtain *CDX2P-CreER^T2^;Apc^15lox/+^* mice, while *Kras^LSL-G12D/+^* mice were crossed with *Trp53^R172H/+^* mice to generate *Kras^LSL-G12D/+^;Trp53^R172H/+^*mice. The resulting strains were interbred to obtain *CDX2P-CreER^T2^;Apc^15lox/+^;Kras^LSL-G12D/+^;Trp53^R172H/+^*mice, hereafter referred to as AKPC mice. *Col1a2-CreER^T2^;Ghr^fl/fl^*mice were generated by crossing *Col1a2-CreER^T2^* mice with *Ghr^flox^*mice.

Cre-mediated recombination in AKPC and *Col1a2-CreER^T2^;Ghr^fl/fl^*mice was induced by intraperitoneal injection of tamoxifen (100 mg/kg body weight; T5648, Sigma-Aldrich, St. Louis, MO, USA) once daily for five consecutive days. Mice were maintained on a mixed C57BL/6J genetic background under specific pathogen-free conditions. Tissues were collected at the indicated time points after tamoxifen administration.

Mice were genotyped at 3 weeks of age using genomic DNA extracted from tail biopsies. Briefly, approximately 3–5 mm of tail tissue was collected and incubated with 50 μL of digestion buffer (SB-201, ShareBio) at 55°C for 15 min in a water bath or heating block. The samples were then heated at 95°C for 5 min to inactivate the protease and centrifuged at 12,000 × g for 5 min. The supernatant was collected and used as the PCR template. PCR amplification was performed in a 25 μL reaction mixture containing 12.5 μL of 2× DetPCR SuperMix (SB-201, ShareBio), 1 μL of gene-specific primers, 1 μL of genomic DNA template, and 10.5 μL of nuclease-free water. The primers used for genotyping are listed in **Table S3**.

For AKPC mice, the *Apc^15lox^*, *Kras^LSL-G12D^*, *Trp53^R172H^*, and *CDX2P-CreER^T2^* alleles were identified by PCR. Expected products were 225 and 380 bp for the wild-type and *Apc^15lox^*alleles, respectively; 250 and 100 bp for the wild-type and *Kras^LSL-G12D^* alleles, respectively; and 297 and 350 bp for the wild-type and *Trp53^R172H^* alleles, respectively. The *CDX2P-CreER^T2^* transgene was identified by a 413-bp product. For *Col1a2-CreER^T2^;Ghr^fl/fl^* mice, the wild-type and targeted *Col1a2-CreER^T2^* alleles yielded 786- and 447-bp products, respectively, whereas the wild-type and floxed *Ghr* alleles yielded 266- and 371-bp products, respectively.

PCR cycling conditions were optimized for each allele. For *Apc^15lox^*, amplification consisted of an initial denaturation at 94°C for 2 min; 5 cycles each of 94°C for 10 s, annealing at 64°C, 61°C, or 58°C for 30 s, and 68°C for 40 s; followed by 35 cycles of 94°C for 10 s, 55°C for 30 s, and 68°C for 40 s; and a final extension at 68°C for 7 min. For *Kras^LSL-G12D^*, amplification consisted of 94°C for 3 min; 10 cycles of 94°C for 20 s, 65°C for 15 s, and 68°C for 10 s; 28 cycles of 94°C for 15 s, 60°C for 15 s, and 72°C for 10 s; and a final extension at 72°C for 2 min. For *Trp53^R172H^*, amplification consisted of 94°C for 3 min, followed by 35 cycles of 94°C for 30 s, 64°C for 1 min, and 72°C for 1 min, with a final extension at 72°C for 2 min. For *CDX2P-CreER^T2^*, amplification consisted of 94°C for 3 min, followed by 35 cycles of 94°C for 30 s, 60°C for 35 s, and 72°C for 35 s, with a final extension at 72°C for 5 min. *Col1a2-CreER^T2^* and *Ghr^flox^* were amplified using an initial denaturation at 95°C for 5 min, followed by 35 cycles of 98°C for 30 s, 58°C for 30 s, and 72°C for 45 s, and a final extension at 72°C for 5 min.

PCR products were separated on 1.5–2.5% agarose gels (SB-P001, ShareBio) prepared in 1× TAE buffer (SB-BR005, ShareBio) and stained with SerRed nucleic acid stain (G3606, Servicebio, Wuhan, China). Electrophoresis was performed at 120–140 V for 30–50 min, depending on gel concentration and expected product size. PCR bands were visualized using a gel imaging system (Tanon).

### Mouse pituitary snRNA-seq and analysis

Nuclei were isolated from frozen pituitary tissues using a Nucleus Isolation Kit (SHBIO 52009-10) and adjusted to a concentration of 700–1,200 nuclei/μL. Single-nucleus RNA-seq libraries were generated using the 10x Genomics Chromium Single Cell 3′ kit (v3) and sequenced on an Illumina NovaSeq platform with paired-end 150-bp reads. Raw sequencing data were processed using Cell Ranger 7.1.0 with the mm10-2020-A mouse reference transcriptome, with intronic reads included in the count matrix. Downstream analyses were performed in Seurat. Nuclei with fewer than 400 detected genes, fewer than 1,000 unique molecular identifiers (UMIs), or more than 20% mitochondrial reads were excluded, and predicted doublets were removed using DoubletFinder. Data were normalized and scaled, followed by PCA and Harmony integration.

UMAP visualization and graph-based clustering were performed using the first 20 Harmony dimensions at a resolution of 0.5.

Cluster-specific marker genes were identified using MAST with a minimum detection fraction of 0.2 and a log_2_ fold-change threshold of 0.25. Cell identities were assigned according to canonical pituitary marker genes reported previously (*16*) and were further supported by GSVA of MSigDB mouse GO biological process gene sets. GSVA was further used to compare pathway activity between AKP and AKPC samples within individual cell types, and differential pathway activity was assessed using limma.

### Plasma proteomics

Peripheral blood was collected from tumor-bearing and control mice at the indicated endpoints, and plasma was isolated by centrifugation and stored at −80°C until analysis. High-abundance plasma proteins were depleted before proteomic sample preparation. Proteins were reduced and alkylated, precipitated, digested with trypsin, and desalted before nano-flow liquid chromatography–mass spectrometry analysis. Peptides were analyzed in data-independent acquisition (DIA) mode, and the resulting data were processed using Spectronaut 18 with directDIA. Precursor and protein identifications were controlled at a 1% false discovery rate (FDR). Protein abundance was quantified using MaxLFQ with cross-run normalization, without missing-value imputation. Differential protein abundance between tumor-bearing and control mice was assessed using an unpaired *t* test with Benjamini–Hochberg correction; proteins with an absolute log_2_ fold change > 0.585 and *q* < 0.05 were considered differentially abundant.

Differentially abundant proteins were subjected to Gene Ontology cellular-component enrichment analysis using the clusterProfiler R package with org.Mm.eg.db as the mouse annotation database. *P* values were adjusted using the Benjamini–Hochberg method.

Upregulated proteins were further intersected with secreted proteins annotated in UniProt/Swiss-Prot using subcellular-localization annotations downloaded in April 2025 (*17*). Candidate secreted proteins with documented receptors were prioritized using CellPhoneDB ligand–receptor annotations together with receptor expression in pituitary bulk RNA-seq and snRNA-seq datasets.

### ELISA assays

Blood samples were collected from mice at the indicated experimental endpoints during a consistent time window. Plasma was separated by centrifugation and stored at −80°C until analysis. Concentrations of GH (EM30221M), prolactin (EM30438M), ACTH (EM30600M), α-MSH (EM30942M), β-endorphin (EM30755M), FSH (EM30583M), LH (EM30672M), and TSH (EM30774M) were measured in plasma or culture supernatants from primary pituitary cells using mouse ELISA kits from Weiao Biotech (Shanghai, China), according to the manufacturer’s instructions. Plasma BMP2 concentrations were measured using a mouse BMP2 ELISA kit (EM30048M, Weiao Biotech). Samples were analyzed within the range of the corresponding standard curves.

### Primary pituitary cultures and treatments

Primary pituitary cells were isolated from female mice as previously described (*5*) with minor modifications. Briefly, freshly isolated pituitary glands were rinsed with calcium- and magnesium-free Hanks’ balanced salt solution (HBSS; 14175095, Gibco), minced, and digested in calcium- and magnesium-containing HBSS (14025092, Gibco) supplemented with 2 mg/mL collagenase II (GC305014, Servicebio), 3% bovine serum albumin (BSA; A500023-0100, Sangon Biotech, China), and 25 mM HEPES (15630080, Gibco) for 1 h at 37°C with gentle shaking. Dispersed cells were washed with calcium- and magnesium-free HBSS and resuspended in DMEM/F12 (11320033, Gibco) supplemented with 15% FBS. Cells were cultured for 1–2 d before treatment.

For immunoblotting, primary pituitary cells were seeded in 6-well plates and treated with rmBMP2 at the indicated concentrations for 45 min or with 25 ng/mL rmBMP2 for the indicated time points, followed by analysis of SMAD1/5 signaling. To assess the effect of *Bmpr1a* knockdown on BMP signaling, primary pituitary cells isolated from AAV-shNC- or AAV-sh*Bmpr1a*-treated mice were treated with 25 ng/mL rmBMP2 for 45 min, followed by analysis of SMAD1/5 signaling. For qRT-PCR, cells were seeded in 24-well plates and treated with 25 ng/mL rmBMP2 for 1 or 6 h, followed by analysis of the BMP-responsive genes *Id1, Id2, Id3,* and *Smad6*. For hormone-secretion assays, pooled primary pituitary cells were evenly distributed into 48-well plates, with each well containing cells corresponding to approximately three pituitaries in 250 μL culture medium. Cells were treated with 25 ng/mL rmBMP2 or vehicle for 24, 48, or 72 h, and adenohypophyseal hormone concentrations in the culture supernatants were measured by ELISA as described above.

### In vivo modulation of BMP signaling

#### Recombinant BMP2 treatment

To test whether BMP2 was sufficient to activate pituitary endocrine output, 8-week-old female C57BL/6J mice received intraperitoneal recombinant mouse BMP2 (10 μg/kg; HY-P7006, MedChemExpress) or vehicle once daily for three consecutive days, after which plasma pituitary hormone concentrations were measured as described above.

#### Recombinant Noggin treatment

To inhibit BMP signaling, mice bearing orthotopic MC38 tumors were intraperitoneally administered recombinant mouse Noggin (25 μg/kg; HY-P7086, MedChemExpress) or an equal volume of vehicle every other day for 3 weeks, starting 1 week after tumor implantation.

#### Stereotaxic virus injection

To suppress *Bmpr1a* expression in the pituitary, AAV2/9-U6-*shBmpr1a* or the scramble control vector AAV2/9-U6-shNC (Obio Technology, Shanghai, China) was delivered bilaterally by stereotaxic injection. Each mouse received approximately 3 × 10^9^ vector genomes (vg) in total, with 500 nL injected per side at a rate of 100 nL/min. The pituitary was targeted using the following coordinates relative to bregma: anteroposterior, −2.5 mm; mediolateral, ±0.15–0.25 mm; and dorsoventral, −6.35 to −6.45 mm. The *Bmpr1a*-targeting shRNA sequence was 5′-GGGTCGTTACAACCGTGATTT-3′. Mice were anesthetized with pentobarbital (50 mg/kg, i.p.) during surgery, and body temperature was maintained at 36–37°C. Orthotopic MC38 tumor experiments were initiated 2 weeks after viral injection.

### Analysis of public mouse scRNA-seq datasets

#### MC38 tumor scRNA-seq data collection and quality control

Single-cell RNA-sequencing data from MC38 tumors in C57BL/6J mice were collected from four publicly available GEO datasets (GSE220635, GSE245465, GSE247841, and GSE263524), comprising seven samples. Only samples from the corresponding control conditions in each study were included. Data were analyzed in R (v4.3.3) using Seurat (v5.2.1). For each sample, cells with ≤400 detected genes were initially excluded, and potential doublets were identified and removed using DoubletFinder (v2.0.6). After merging, cells with ≤400 detected genes, ≤1,000 UMIs, or ≥20% mitochondrial transcripts were excluded. A total of 33,474 cells were retained for downstream integrated analysis.

#### MC38 tumor data integration, clustering, and cell-type identification

Gene-expression matrices were normalized using the LogNormalize method, and 2,000 highly variable genes were identified using the variance-stabilizing transformation method. Data were scaled and subjected to PCA. Batch effects among samples were corrected using Harmony (v1.2.3), with sample identity used as the integration variable. The first 20 Harmony dimensions were used to construct the nearest-neighbor graph, perform unsupervised clustering at a resolution of 1.0, and generate UMAP embeddings. Cluster-specific marker genes were identified using MAST (v1.28.0), with a minimum detection fraction of 0.2 and a log_2_ fold-change threshold of 0.25. Major cell types were manually annotated based on the expression of canonical marker genes and included fibroblasts, macrophages, monocytes, cancer cells, CD4^+^ and CD8^+^ T cells, NK cells, dendritic cells, and endothelial cells.

#### Fibroblast-specific analysis

Fibroblasts were extracted from the integrated dataset and independently reprocessed by normalization, identification of 2,000 highly variable genes, scaling, PCA, and Harmony integration using sample identity as the batch variable. Graph-based clustering and UMAP were performed using the first 20 Harmony dimensions at a clustering resolution of 0.5. Fibroblast subclusters were subsequently classified according to their activation state based on the coordinated expression of *Pdgfra*, *Acta2*, *Fap*, *Lrrc15*, and *Pdpn* (*66*). Subclusters exhibiting a prominent activation-associated expression pattern were annotated as activated fibroblasts, whereas the remaining fibroblast subclusters were operationally annotated as resting fibroblasts. For sample-level comparison of *Cdkn1a* and *Cdkn2a*, all fibroblasts from each of the seven independent MC38 tumor samples were analyzed. For each gene, the mean normalized expression level was calculated across all fibroblasts in each sample, and the percentage of fibroblasts with detectable expression was calculated as the proportion of cells with a normalized expression value greater than zero.

#### Analysis of scRNA-seq datasets from peripheral immune and hematopoietic tissues

Publicly available scRNA-seq datasets from mouse lymph nodes, bone marrow, and spleen were obtained from GEO. Lymph node samples (GSM8496432 and GSM8496433), bone marrow samples (GSM8496449 and GSM8496450), and an independent spleen sample (GSM8496446) were obtained from GSE276312. Additional spleen samples (GSM6661396 and GSM6661398) were obtained from GSE216189. Only control or untreated wild-type samples were included in the analysis.

Raw 10X count matrices were processed using Seurat. Genes detected in fewer than three cells were excluded during Seurat object construction. Cells with ≤200 or ≥6,000 detected genes or with ≥10% mitochondrial transcripts were removed. Data were normalized using the LogNormalize method with a scale factor of 10,000, and the 2,000 most variable genes were identified using the variance-stabilizing transformation method. Data were scaled and subjected to principal component analysis. The first 20 principal components were used for nearest-neighbor graph construction, graph-based clustering, and UMAP visualization. Cell populations were manually annotated according to canonical lineage markers.

### Analysis of public human CRC transcriptomic datasets

RNA-seq data and patient information for primary CRC samples from The Cancer Genome Atlas colon adenocarcinoma (COAD) and rectal adenocarcinoma (READ) cohorts were obtained from the COADREAD PanCancer Atlas dataset in cBioPortal (https://www.cbioportal.org/study/summary?id=coadread_tcga_pan_can_atlas_2018). The downloaded dataset included batch-corrected RNA-seq by expectation maximization (RSEM)–normalized expression values and corresponding patient information. Consensus molecular subtype (CMS) labels from the original CMS classification study were obtained from Synapse (https://www.synapse.org/#!Synapse:syn2623706/wiki/67246) (*23*). To visualize *GHR* expression across CMS subtypes, RSEM–normalized counts were transformed as log_2_(count + pseudocount), with pseudocount adjustment applied to normalize the lowest *GHR* expression value to zero (**Figure S6C**). Pearson correlations between *GHR* expression and fibroblast activation markers (*ACTA2, FAP, PDGFA, PDPN,* and *LRRC15*) were calculated from the RSEM-normalized expression data.

Associations between *GHR* expression and stromal score or estimated CAF abundance were assessed using TIMER3.0 with the ESTIMATE, EPIC, MCP-counter, xCell, and TIDE algorithms. Survival analyses were performed using the Kaplan–Meier Plotter platform (https://kmplot.com). Overall survival was analyzed according to *GHR* expression, whereas both overall survival and relapse-free survival were analyzed according to *CDKN1A* expression in the overall colon cancer cohort and the CMS4 subgroup.

### In vivo GH administration

To evaluate the role of GH in tumor growth, tumor-bearing mice received subcutaneous GH (1 mg/kg; HY-P70261, MedChemExpress) or vehicle every other day, beginning on day 2 after tumor inoculation and continuing until the end of the experiment.

### Proliferation assay *in vitro*

Cell viability was assessed using a Cell Counting Kit-8 (CCK-8; C6005, NCM Biotech) according to the manufacturer’s instructions. MC38 cells were seeded in 96-well plates at a density of 1–5 × 10^3^ cells per well and subjected to the indicated treatments. Cells were cultured at 37°C in a humidified incubator containing 5% CO_2_. At the indicated time points, CCK-8 reagent was added to each well, and the absorbance at 450 nm (OD_450_) was measured using a Synergy Neo2 multimode microplate reader (BioTek, USA). Cell viability (%) was calculated as (ODobserved − ODblank) / (ODcontrol − ODblank) × 100%.

### Primary CAF isolation and culture

NFs and CAFs were isolated from normal colorectal tissues and CRC tissues of mice, respectively, as previously described with minor modifications (*67*). Briefly, for NF isolation, normal colorectal tissues were placed in culture dishes and compressed beneath a 20-mm coverslip to prevent the tissue from floating. Tissues were cultured in DMEM supplemented with 10% FBS and 1% penicillin–streptomycin, with the medium replaced daily. After 7–10 d, fibroblasts migrated out from the tissue explants and expanded on the culture surface. The residual tissue was then removed, and the fibroblasts were further expanded. For CAF isolation, freshly dissected CRC tissues were rinsed with cold PBS to remove residual blood and debris and minced into small fragments (approximately 1–2 mm^3^). Tissue fragments were digested with collagenase II (GC305014, Servicebio) at 37°C for 1–2 h with gentle agitation until most of the tissue was dissociated. The resulting suspension was passed through a 70-μm cell strainer to remove undigested tissue and subjected to sequential centrifugation to remove residual enzyme and debris. The final cell pellet was resuspended in DMEM supplemented with 10% FBS and 1% penicillin–streptomycin and seeded into culture dishes. After 30 min, adherent fibroblasts were retained, whereas non-adherent cells were removed.

### CAF RNA sequencing

CAF-1 cells were treated with vehicle or GH (500 ng/mL) for 7 d before RNA isolation. Poly(A)-enriched mRNA libraries were prepared using the VAHTS Universal V10 RNA-seq LibraryPrep Kit (Vazyme) and subjected to paired-end sequencing on a DNBSEQ-T7 sequencing platform (MGI). Raw reads were processed using fastp and aligned to the mouse reference genome GRCm39 (Ensembl release 111) using HISAT2. Gene expression was quantified using StringTie and normalized using TMM. Expression levels were represented as FPKM, and differential gene-expression analysis was performed using edgeR. PCA was performed in R on Hellinger-transformed gene-expression data using prcomp and visualized using scatterplot3d (v0.3-44). Heatmaps of representative SASP genes were generated in R using pheatmap, with expression values standardized across samples for each gene by row-wise scaling. GSEA (v4.3.3) was performed to evaluate senescence- and SASP-associated transcriptional programs using the SenMayo gene set (SAUL_SEN_MAYO (*68*)) from MSigDB and a previously published SASP gene signature (*69*). Gene sets with an FDR *q* value < 0.05 were considered significantly enriched. Sequencing data have been deposited in the NCBI BioProject database under accession number PRJNA1512966.

### SA-β-gal staining assay

Indicated CAFs were fixed in 4% formaldehyde at room temperature for 20 min and incubated with freshly prepared 1× SA-β-gal detection solution (C0602, Beyotime) at 37°C for 5 h. SA-β-gal–positive cells were quantified in randomly selected fields using ImageJ software.

### JAK2–STAT5 pathway inhibition

To assess the involvement of JAK2–STAT5 signaling in GH-induced responses, CAF-1 cells were pretreated for 30 min with the JAK2 inhibitor fedratinib (HY-10409, MedChemExpress) or the STAT5 inhibitor STAT5-IN-1 (HY-101853, MedChemExpress), followed by GH stimulation. For immunoblot analysis, the inhibitors were used at the indicated concentrations.

For qRT-PCR and SA-β-gal assays, fedratinib and STAT5-IN-1 were used at 2 μM and 50 μM, respectively. For the 7-day SA-β-gal assay, inhibitors were maintained throughout GH treatment. p21 protein expression, *Cdkn1a* mRNA levels, and SA-β-gal activity were assessed as described above.

### Rectal submucosal AAV injection

For rectal submucosal viral injection, *Col1a2-CreER^T2^* or *Ghr^fl/fl^;Col1a2-CreER^T2^* mice received AAV9-DIO-miR30-*shCdkn1a*, AAV9-DIO-miR30-*shCdkn2a*, or AAV9-DIO-miR30-shNC (Obio Technology, Shanghai, China) at a dose of 2 × 10^11^ vg per mouse. Under a stereomicroscope, the anus was gently exposed, and 2 μL of viral suspension was injected into the rectal submucosa using a microsyringe at a rate of 10 μL/min. After injection, the needle was kept in place for 1 min before being slowly withdrawn. The shRNA sequences used were 5′-CTATCACTCCAAGCGCAGATT-3′ for *Cdkn1a* and 5′-TCAAGACATCGTGCGATATTT-3′ for *Cdkn2a*. Tamoxifen was subsequently administered to induce Cre-dependent shRNA expression, followed by orthotopic implantation of MC38 cells according to the experimental design.

### Dasatinib plus quercetin treatment

For pharmacological senolytic experiments, orthotopic MC38-bearing C57BL/6J mice were treated with dasatinib (5 mg/kg; HY-10181, MedChemExpress) plus quercetin (50 mg/kg; HY-18085, MedChemExpress) by oral gavage three times per week for 3 weeks. p21^+^ CAFs, p21^+^ immune cells, tumor burden, and survival were assessed. To evaluate potential direct effects on tumor cells, MC38 cells were treated with quercetin (15 μM) together with increasing concentrations of dasatinib (0, 1, 2, 5, 10, or 20 μM), and cell viability was measured at 0, 24, 48, and 72 h using the CCK-8 assay as described above.

### H&E staining and immunohistochemistry

Mouse colorectal tumor tissues were fixed overnight in 4% paraformaldehyde, prepared as Swiss rolls, paraffin-embedded, and sectioned at 5 μm. Paraffin sections were deparaffinized, rehydrated, and stained with hematoxylin and eosin using an H&E staining kit (G1076, Servicebio), followed by dehydration, clearing, and mounting. For quantitative assessment of tumor burden, tumor regions and total tissue areas were annotated on whole-section H&E images using QuPath.

For immunohistochemical staining of mouse colorectal tissues, paraffin sections were deparaffinized and rehydrated, followed by heat-induced antigen retrieval in citrate buffer (pH 6.0; G1202, Servicebio). Endogenous peroxidase activity was blocked with 3% hydrogen peroxide for 25 min at room temperature, followed by blocking with 3% BSA for 30 min.

Sections were incubated overnight at 4°C with primary antibodies against Ki67 (GB151499, Servicebio; 1:500), CK20 (GB112050, Servicebio; 1:1,500), or β-catenin (GB150016, Servicebio; 1:1,000). Sections were subsequently incubated with species-appropriate horseradish peroxidase (HRP)-conjugated secondary antibodies (ab6721 or ab6789, Abcam) for 50 min at room temperature. Immunoreactivity was visualized using 3,3′-diaminobenzidine (DAB) substrate (K5007, DAKO), followed by hematoxylin counterstaining, dehydration, clearing, and mounting.

Human CRC formalin-fixed, paraffin-embedded (FFPE) sections were subjected to immunohistochemical staining using the same procedure and incubated with an anti-p21 primary antibody (HA601018, HUABIO; 1:400) overnight at 4°C. For quantitative analysis of p21 staining, stromal/epithelial regions were manually annotated as regions of interest in QuPath.

Cells were detected based on hematoxylin-stained nuclei, and p21-positive nuclei were identified based on nuclear DAB staining using a fixed threshold applied uniformly across all samples. For each sample, 3–5 randomly selected fields were analyzed.

### Immunofluorescence

For immunofluorescence staining of primary CAFs, cells were seeded in chamber slides (ibidi), washed with PBS, and fixed with 4% paraformaldehyde for 15 min at room temperature. Cells were permeabilized with 0.5% Triton X-100 for 10 min at room temperature and blocked with 10% BSA for 1 h. Primary antibodies, including α-SMA (14395-1-AP, Proteintech; 1:800), FAP (AF6858, Beyotime; 1:200), and GHR (bs-0654R, Bioss; 1:500), were diluted in 1% BSA and incubated with the cells overnight at 4°C. After washing, cells were incubated with appropriate fluorophore-conjugated secondary antibodies diluted in 1% BSA for 1 h at room temperature in the dark. Nuclei were counterstained with DAPI for 10 min at room temperature.

For multiplexed immunofluorescence, tissues were fixed in 4% paraformaldehyde for 24 h, paraffin-embedded, and sectioned at 4 μm. Sections were deparaffinized, rehydrated through graded ethanol, and subjected to antigen retrieval. After blocking endogenous peroxidase activity and nonspecific binding, tissue sections underwent iterative staining cycles comprising: (1) overnight incubation with primary antibodies at 4°C, (2) incubation with species-appropriate HRP-conjugated secondary antibodies for 50 min at room temperature, (3) tyramide signal amplification (TSA) using distinct fluorophores for 10 min at room temperature, and (4) antibody stripping to remove bound primary and secondary antibodies before the subsequent staining cycle. Primary antibodies included α-SMA (ab124964, Abcam; 1:500), p21 (ab188224, Abcam; 1:500), and CD45 (HA723606, HUABIO; 1:500). Following completion of all TSA cycles, nuclei were counterstained with DAPI for 10 min at room temperature, tissue autofluorescence was quenched, and sections were mounted with antifade mounting medium.

All fluorescence images were acquired using a confocal microscope (Leica, Germany). Quantitative image analysis was performed using QuPath (v0.7.0). Tumor regions were randomly selected, with a total analyzed area of at least 2 mm^2^ per mouse. Total cell numbers in analyzed images were scored using the “cell detection” command based on the DAPI channel. Nuclear p21 positivity was determined using the mean nuclear fluorescence intensity (Nucleus: Mean), with a fixed positivity threshold applied uniformly across all samples. α-SMA-positive and CD45-positive cells were defined based on their respective cellular fluorescence intensities (Cell: Mean) using fixed thresholds that were consistently applied across samples.

### Quantitative reverse-transcription PCR (qRT-PCR)

Total RNA was extracted from frozen tissues or cultured cells using TRIzol reagent (15596018, Thermo Fisher Scientific) and reverse-transcribed into cDNA using the All-in-One First-Strand Synthesis MasterMix with dsDNase (SB-RT001, ShareBio) according to the manufacturer’s instructions. The resulting cDNA was diluted 1:6 and subjected to qRT-PCR using SYBR Green qPCR Premix (SB-Q204, ShareBio) on a QuantStudio 3 Real-Time PCR System (Applied Biosystems). Relative gene expression was calculated using the 2^−ΔΔCt^ method and normalized to *Gapdh*. Primer sequences used for qRT-PCR are listed in **Table S3**.

### SDS-PAGE and immunoblot analysis

Cells were washed with ice-cold PBS and lysed on ice for 20 min in RIPA lysis buffer (50 mM Tris-HCl, pH 7.6, 150 mM NaCl, 1% Triton X-100, 0.5% sodium deoxycholate, and 0.1% SDS; WB3100, NCM Biotech) supplemented with protease and phosphatase inhibitor cocktail (P002, NCM Biotech). Lysates were clarified by centrifugation at 12,000 × *g* for 15 min at 4°C, and protein concentrations were determined using a BCA protein assay kit (P0011, Beyotime).

Protein samples were mixed with SDS-PAGE loading buffer (WB2001, NCM Biotech), heated at 95–100°C for 10 min, and separated by SDS-PAGE using 7.5–12.5% resolving gels (NCM Biotech). Proteins were transferred onto 0.2- or 0.45-μm nitrocellulose membranes (10600001 and 10600002, Cytiva) at 400 mA for 25–40 min using a Bio-Rad transfer system. Membranes were blocked for 25 min at room temperature in TBST containing 5% nonfat milk or, for phosphoprotein detection, 5% BSA, and incubated with the indicated primary antibodies at 4°C overnight. After washing, membranes were incubated with HRP-conjugated secondary antibodies (SA00001-1 or SA00001-2, Proteintech) for 1 h at room temperature. Protein signals were detected using enhanced chemiluminescence substrate (P2300, NCM Biotech) and imaged using a Tanon 5200 imaging system.

Primary antibodies included GAPDH (60004-1-Ig, Proteintech; 1:5,000), phospho-SMAD1/5/9 (13820, Cell Signaling Technology; 1:1,000), SMAD1 (10429-1-AP, Proteintech; 1:1,000), SMAD5 (12167-1-AP, Proteintech; 1:2,000), FAP (AF6858, Beyotime; 1:1,000), α-SMA (14395-1-AP, Proteintech; 1:2,000), p21 (28248-1-AP, Proteintech; 1:2,000), and p16INK4a (YM8152, Immunoway; 1:2,000).

### PET/CT imaging and pituitary SUVR analysis

Patients were required to fast for 6 h before PET/CT scanning. ^18^F-FDG was administered intravenously, and PET/CT imaging was performed 60 min after injection using a uMI S-96R PET/CT scanner (United Imaging Healthcare, Shanghai, China). PET/CT images were viewed and analyzed using Bee DICOM Viewer (v3.7.2; SinoUnion Healthcare Inc.).

For quantification of pituitary ^18^F-FDG uptake, the pituitary gland was identified with anatomical guidance from the corresponding CT images. A region of interest (ROI) was manually delineated along the boundary of the entire pituitary gland on the axial slice showing its largest cross-sectional area, and the mean standardized uptake value (SUVmean) within the ROI was recorded. A reference ROI was manually delineated in the cerebellar gray matter, and its SUVmean was measured. Pituitary standardized uptake value ratio (SUVR) was calculated as pituitary SUVmean / cerebellar gray matter SUVmean. ROI delineation was performed by one nuclear medicine physician and reviewed by a second nuclear medicine physician. PET/CT images acquired between September 2024 and June 2026 were included. Patients with known pituitary or sellar lesions, major intracranial structural abnormalities, or a history of intracranial surgery or radiotherapy were excluded. For patients with CRC, only patients with active CRC at the time of PET/CT imaging were included. For analyses involving tumor burden, only patients with newly diagnosed CRC who had not received prior antitumor treatment were included.

For primary CRC lesions, ROIs were manually delineated over the tumor on PET/CT images, and the maximum standardized uptake value (SUVmax) was automatically calculated using the image-analysis software. Tumor dimensions were obtained from the corresponding pathological records, and tumor volume was estimated as length × width × height / 2 (cm^3^).

### Statistical analysis

Data are presented as mean ± SEM unless otherwise indicated. Statistical analyses were performed using GraphPad Prism (v10.1.2) and R (v4.3.3). Comparisons between two independent groups were performed using two-tailed unpaired Student’s *t* tests unless otherwise specified. Paired nonparametric comparisons were performed using two-tailed paired Wilcoxon signed-rank tests. Comparisons among three or more groups were performed using one-way ANOVA with Tukey’s multiple comparisons test. Correlations between continuous variables were assessed using Pearson or Spearman correlation coefficients, as indicated in the corresponding figure legends. Survival curves were estimated using the Kaplan–Meier method and compared using the log-rank (Mantel–Cox) test. Multiple-testing correction for transcriptomic and proteomic analyses was performed as described in the corresponding sections. All statistical tests were two-sided unless otherwise specified, and *P* < 0.05 was considered statistically significant.

The sample size for *in vivo* and *in vitro* experiments was based on previous experience. Where applicable, animals were randomly assigned to treatment groups, and investigators were blinded to group allocation during outcome assessment. All statistical analyses were performed using at least three biological or independent replicates. *n* represents biologically independent samples or animals unless otherwise indicated. Sample sizes are provided in the corresponding figure legends. No samples or animals were excluded from the analyses.

## Acknowledgments

We thank Y. Hao for providing mouse strains.

## Author contributions

W.L., M.Y., S.J., and J.X. conceived the study, designed and performed the experiments, interpreted the results, and wrote the manuscript. S.W., H. Wu, and C.S. performed experiments. S.L. and H. Wang performed computational and statistical analyses. R.H., K.H., and C.Z. acquired clinical data.

## Data, code, and materials availability

All data are available in the main text or the supplementary materials.

---

**Fig. S1.**
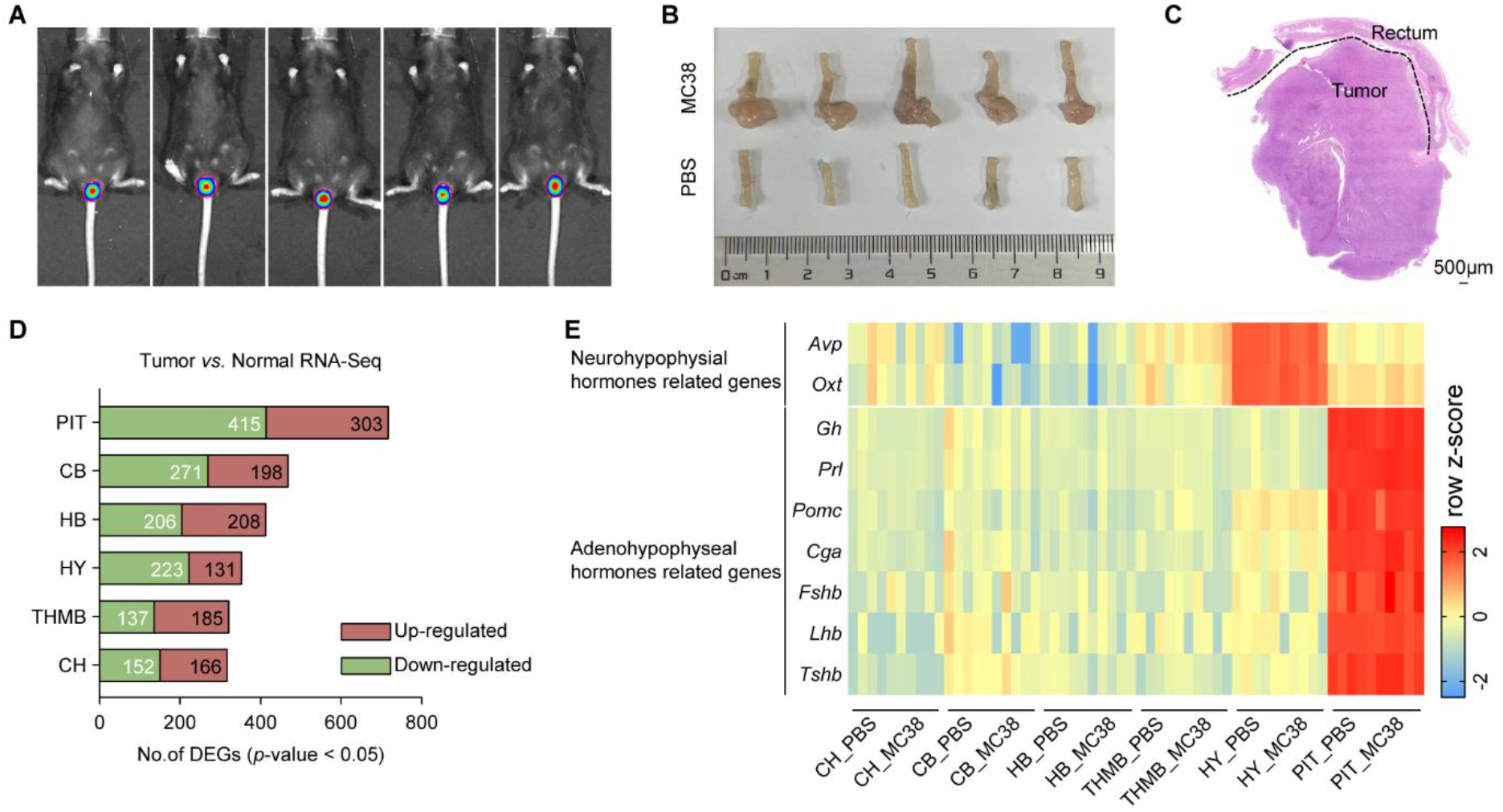
Establishment of an orthotopic MC38 CRC model and transcriptomic analysis of the brain and pituitary. (**A**) In vivo imaging analysis of tumor progression at day 14 after MC38 inoculation. (**B**) Gross morphology of rectal tumors in mice at 28 d post tumor cell injection. (**C**) Representative H&E staining image of rectal tumors at 28 d post tumor cell injection. (**D**) Summary of differentially expressed genes (DEGs) in dissected brain regions and the pituitary gland. CH, cerebrum; CB, cerebellum; HB, hindbrain; THMB, thalamus with midbrain; HY, hypothalamus; PIT, pituitary. (**E**) Heatmap of neurohypophysial- and adenohypophyseal hormone–related genes across dissected brain regions in control and MC38 tumor–bearing mice, measured by RNA-seq, with data transformed into z scores.

**Fig. S2.**
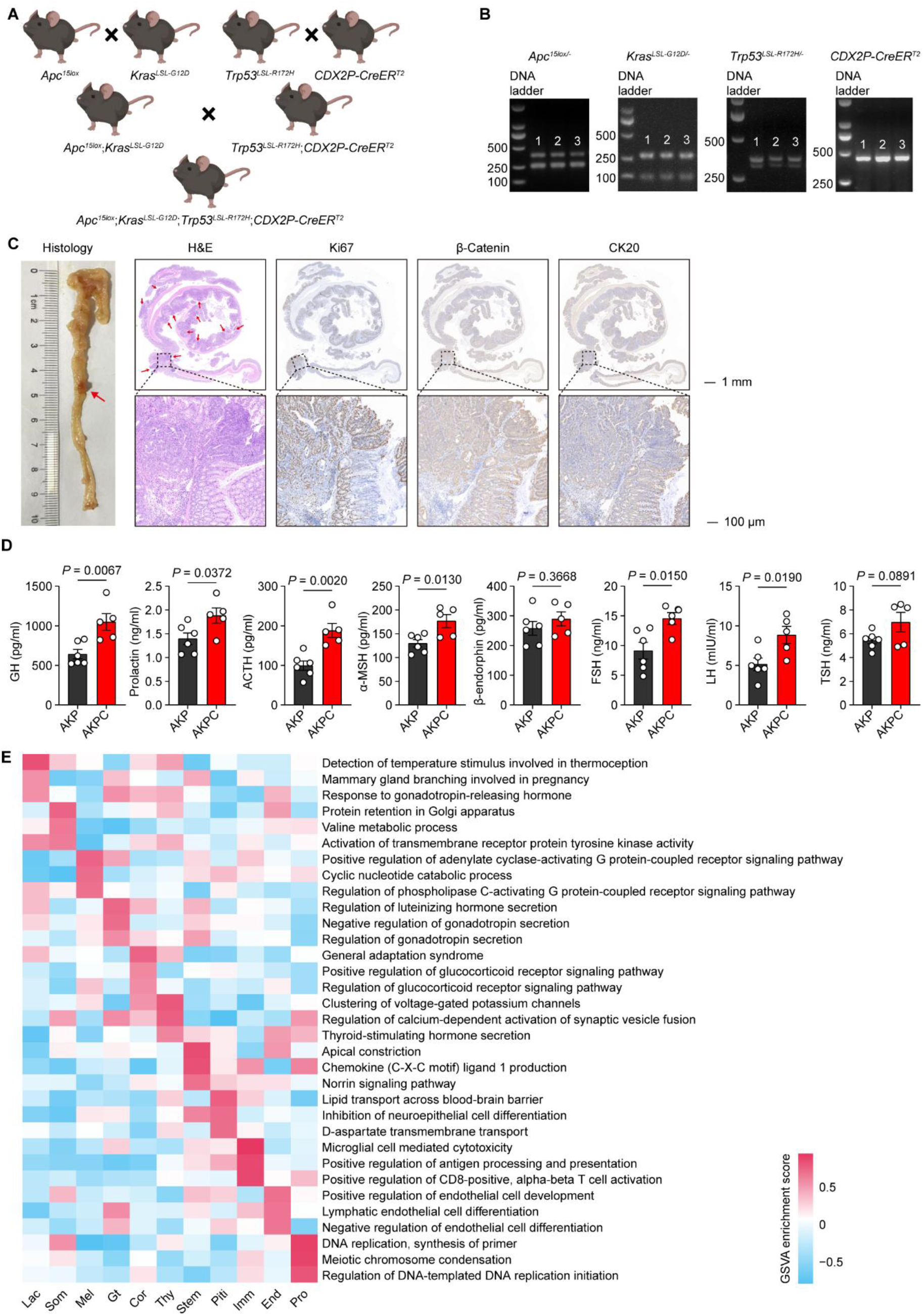
Generation and phenotypic characterization of AKPC mice and associated pituitary endocrine changes. (**A** and **B**) Schematic and gel verification of AKPC mouse generation. (**C**) Representative gross photograph, hematoxylin and eosin (H&E) staining, and immunohistochemical (IHC) staining of Ki67, β-catenin, and CK20 in colorectal tissues from AKPC mice. (**D**) ELISA analysis of plasma adenohypophyseal hormones in AKP and AKPC mice at 8 weeks after tamoxifen induction. AKP, *n* = 6 mice; AKPC, *n* = 5 mice. GH, growth hormone; prolactin (PRL); ACTH, adrenocorticotropic hormone; α-MSH, α-melanocyte-stimulating hormone; β-endorphin; FSH, follicle-stimulating hormone; LH, luteinizing hormone; TSH, thyroid-stimulating hormone. (**E**) Heatmap showing different pathways enriched in integrated pituitary cell clusters from snRNA-seq data by GSVA analysis, colored by mean GSVA enrichment scores. The data are presented as mean ± SEM. *P* values are determined by two-tailed unpaired Student’s *t* test (D). Figure A created with BioRender.com.

**Fig. S3.**
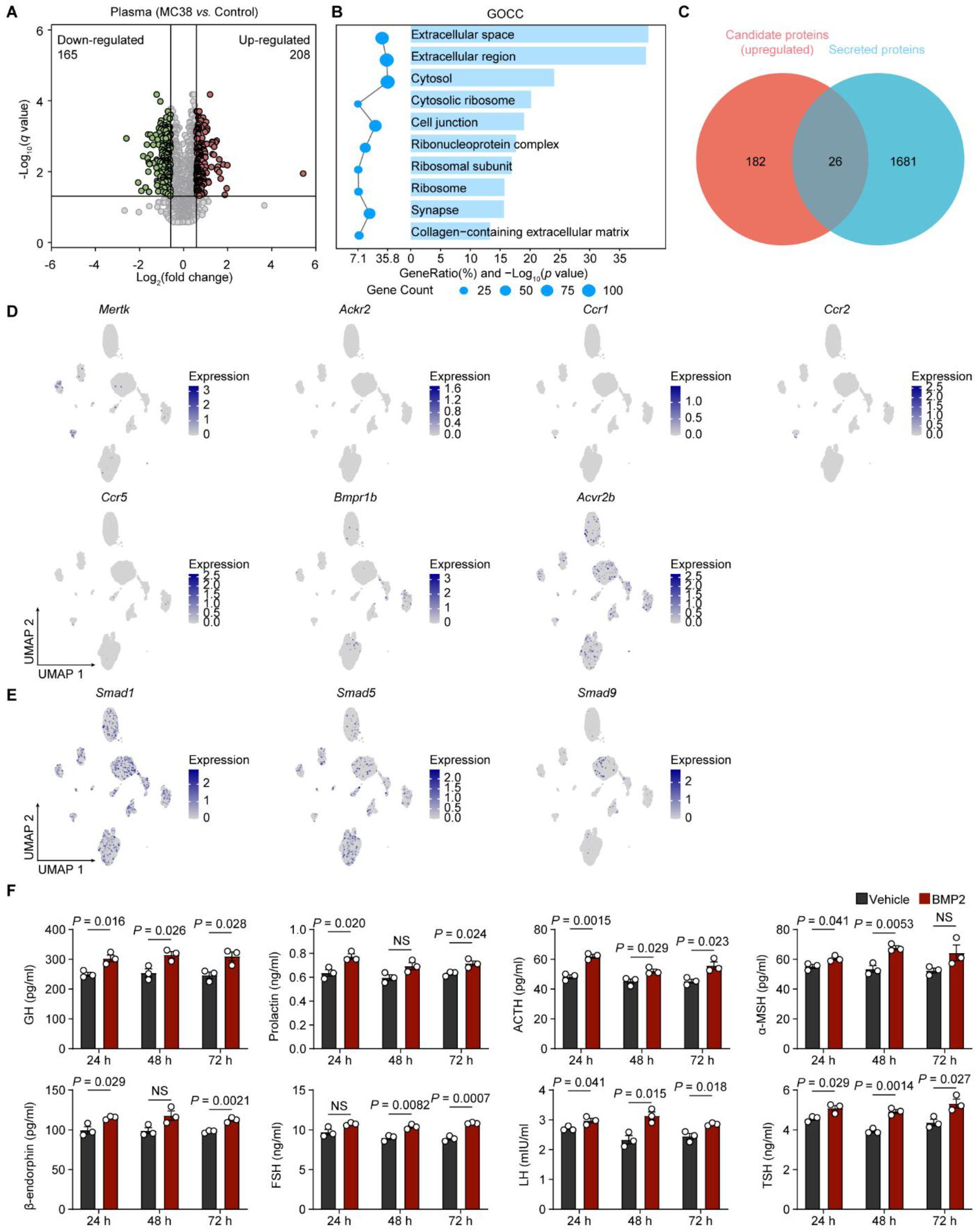
CRC plasma proteome analysis and *in vitro* functional validation of BMP2 in primary pituitary cells. (**A** to **C**) Proteomics analysis of plasma proteins isolated from MC38 tumor-bearing mice and tumor-free control mice. (A) Volcano plot of differential plasma proteins between MC38 tumor-bearing mice and tumor-free control mice. See also Table S1. (B) Bubble plot showing top enriched Gene Ontology Cellular Component (GOCC) terms of differentially expressed plasma proteins from MC38 tumor-bearing versus control mice. Differential proteins were filtered with thresholds of |log_2_(fold change)| > 0.585 and *q* < 0.05. (C) Venn diagram showing overlapping proteins between tumor-induced upregulated candidate plasma proteins and known secreted proteins. (**D** and **E**) snRNA-seq UMAP feature plots showing pituitary cell expression patterns of candidate protein receptors (D) and SMAD family transcription factors (E). (**F**) ELISA analysis of adenohypophyseal hormones in culture supernatants from primary mouse pituitary cells treated with rmBMP2 (25 ng/mL) or vehicle for 24, 48, and 72 h. GH, growth hormone; PRL, prolactin; ACTH, adrenocorticotropic hormone; α-MSH, α-melanocyte-stimulating hormone; FSH, follicle-stimulating hormone; LH, luteinizing hormone; TSH, thyroid-stimulating hormone (*n* = 3). The data are presented as mean ± SEM. Comparisons between vehicle- and rmBMP2-treated cells at each time point were performed using a two-tailed unpaired Student’s *t* test (F). NS, not significant.

**Fig. S4.**
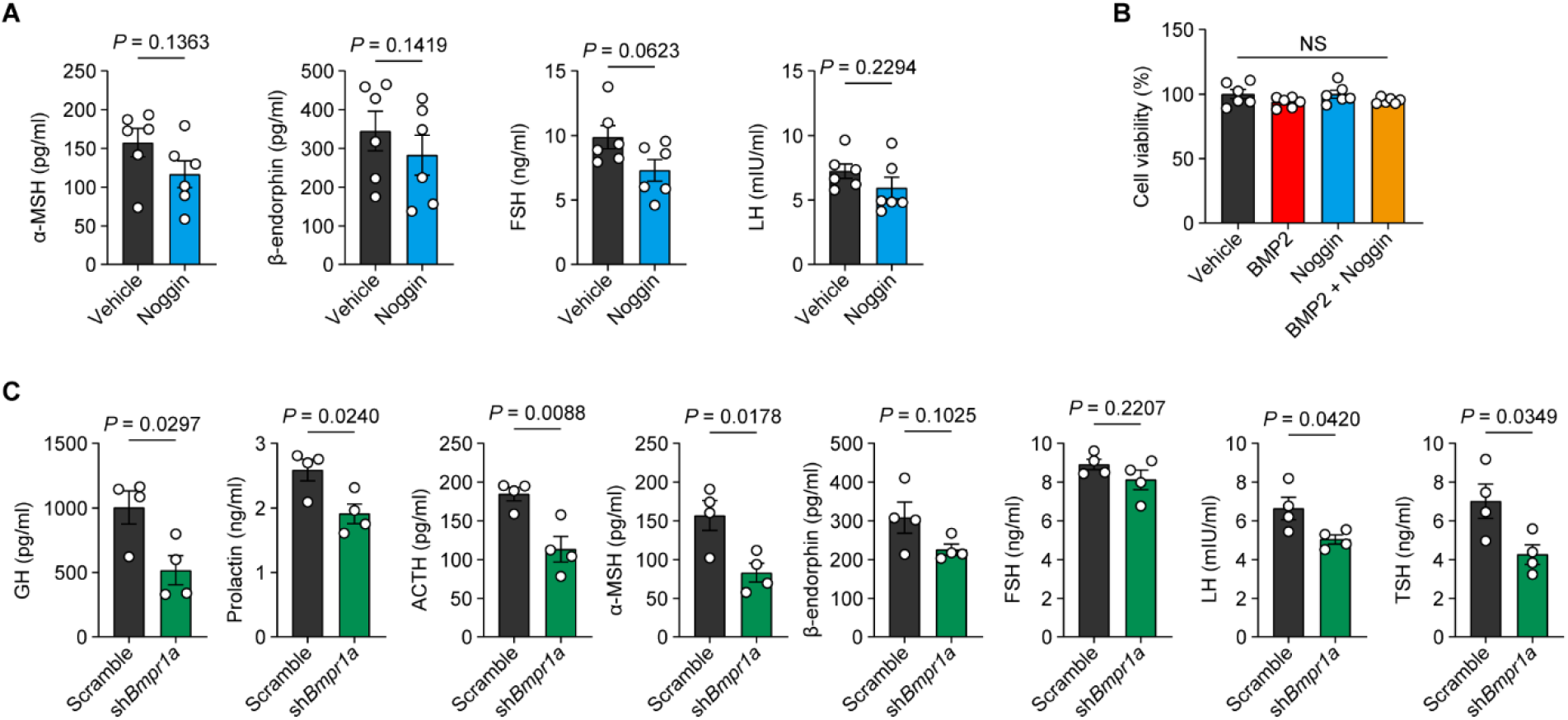
Pituitary BMP signaling regulates endocrine output without directly affecting MC38 cell viability. (**A**) ELISA quantification of plasma pituitary hormones including α-MSH, β-endorphin, FSH and LH in vehicle- and Noggin-treated mice (*n* = 6). (**B**) Cell viability of MC38 cells treated with vehicle, rmBMP2 (25 ng/mL), rmNoggin (100 ng/mL), or rmBMP2 plus rmNoggin for 48 h (n = 6 biological replicates per group). (**C**) ELISA quantification of plasma pituitary hormones including GH, prolactin, ACTH, α-MSH, β-endorphin, FSH, LH and TSH in scramble- and sh*Bmpr1a*-treated mice (*n* = 4). The data are presented as mean ± SEM. *P* values are determined by two-tailed unpaired Student’s *t* test (A and C) and one-way ANOVA with Tukey’s multiple comparisons test (B). NS, not significant.

**Fig. S5.**
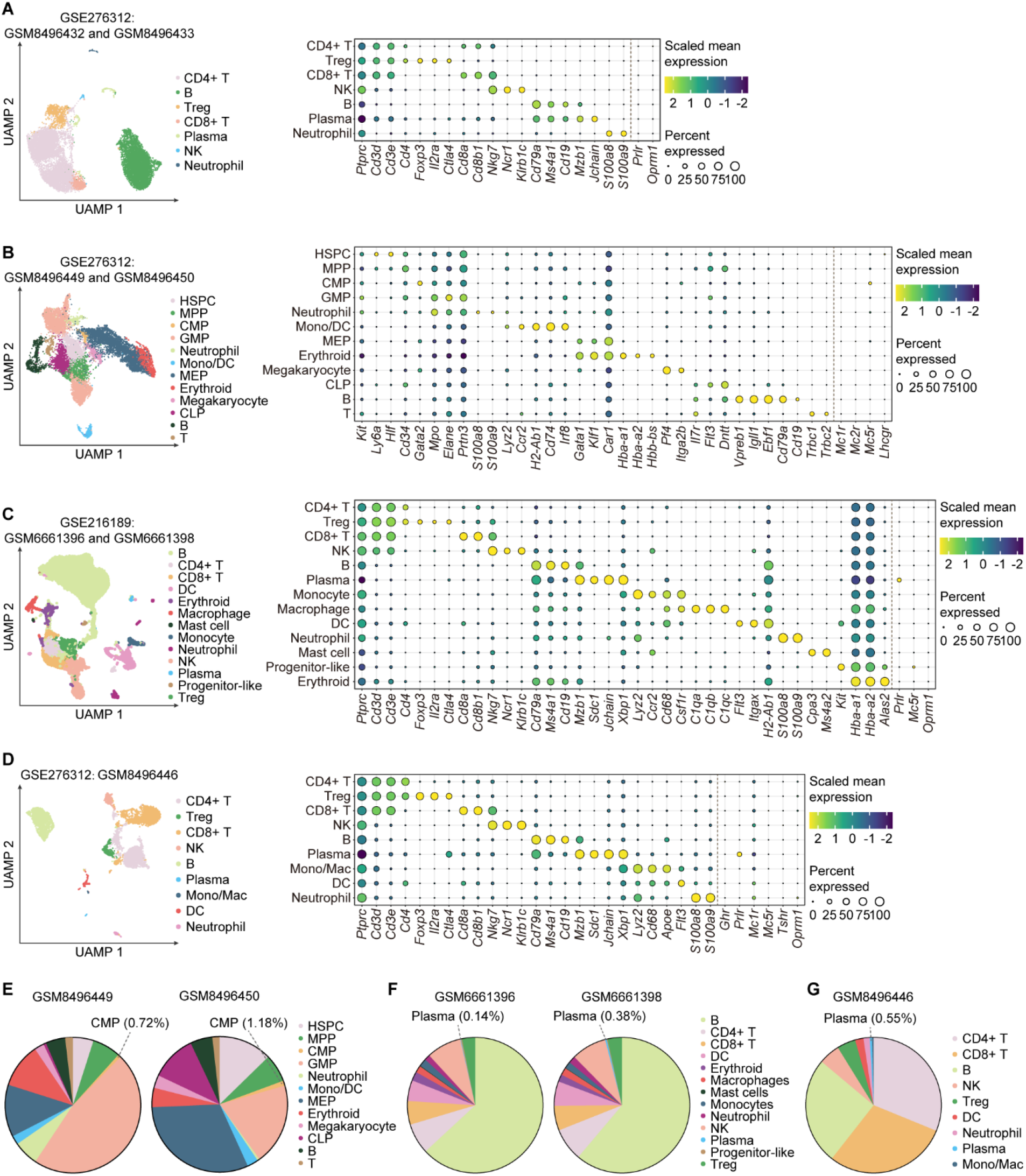
Single-cell survey of pituitary hormone receptor expression in peripheral immune and hematopoietic tissues. (**A**) Single-cell RNA-seq analysis of mouse lymph nodes from GSE276312 (GSM8496432 and GSM8496433). Left, UMAP visualization of major immune cell populations. Right, dot plot showing representative cell-type marker genes and pituitary hormone receptor expression across annotated cell populations. (**B**) Single-cell RNA-seq analysis of mouse bone marrow from GSE276312 (GSM8496449 and GSM8496450). Left, UMAP visualization of major hematopoietic and immune cell populations. Right, dot plot showing representative cell-type marker genes and pituitary hormone receptor expression across annotated cell populations. (**C**) Single-cell RNA-seq analysis of mouse spleen samples from GSE216189 (GSM6661396 and GSM6661398). Left, UMAP visualization of major splenic cell populations. Right, dot plot showing representative cell-type marker genes and pituitary hormone receptor expression. (**D**) Single-cell RNA-seq analysis of an independent mouse spleen dataset from GSE276312 (GSM8496446). Left, UMAP visualization of major immune cell populations. Right, dot plot showing representative cell-type marker genes and pituitary hormone receptor expression. (**E**) Cellular composition of the two bone marrow samples analyzed in (B). The proportion of CMPs among all analyzed cells is indicated for each sample. (**F**) Cellular composition of the two spleen samples analyzed in (C). The proportion of plasma cells among all analyzed cells is indicated for each sample. (**G**) Cellular composition of the spleen sample analyzed in (D). The proportion of plasma cells among all analyzed cells is indicated. In the dot plots, pituitary hormone receptors with no detectable expression in a given dataset were not shown. HSPC, hematopoietic stem and progenitor cell; MPP, multipotent progenitor; CMP, common myeloid progenitor; GMP, granulocyte–monocyte progenitor; MEP, megakaryocyte–erythroid progenitor; CLP, common lymphoid progenitor; Mono, monocyte; Mac, macrophage; DC, dendritic cell.

**Fig. S6.**
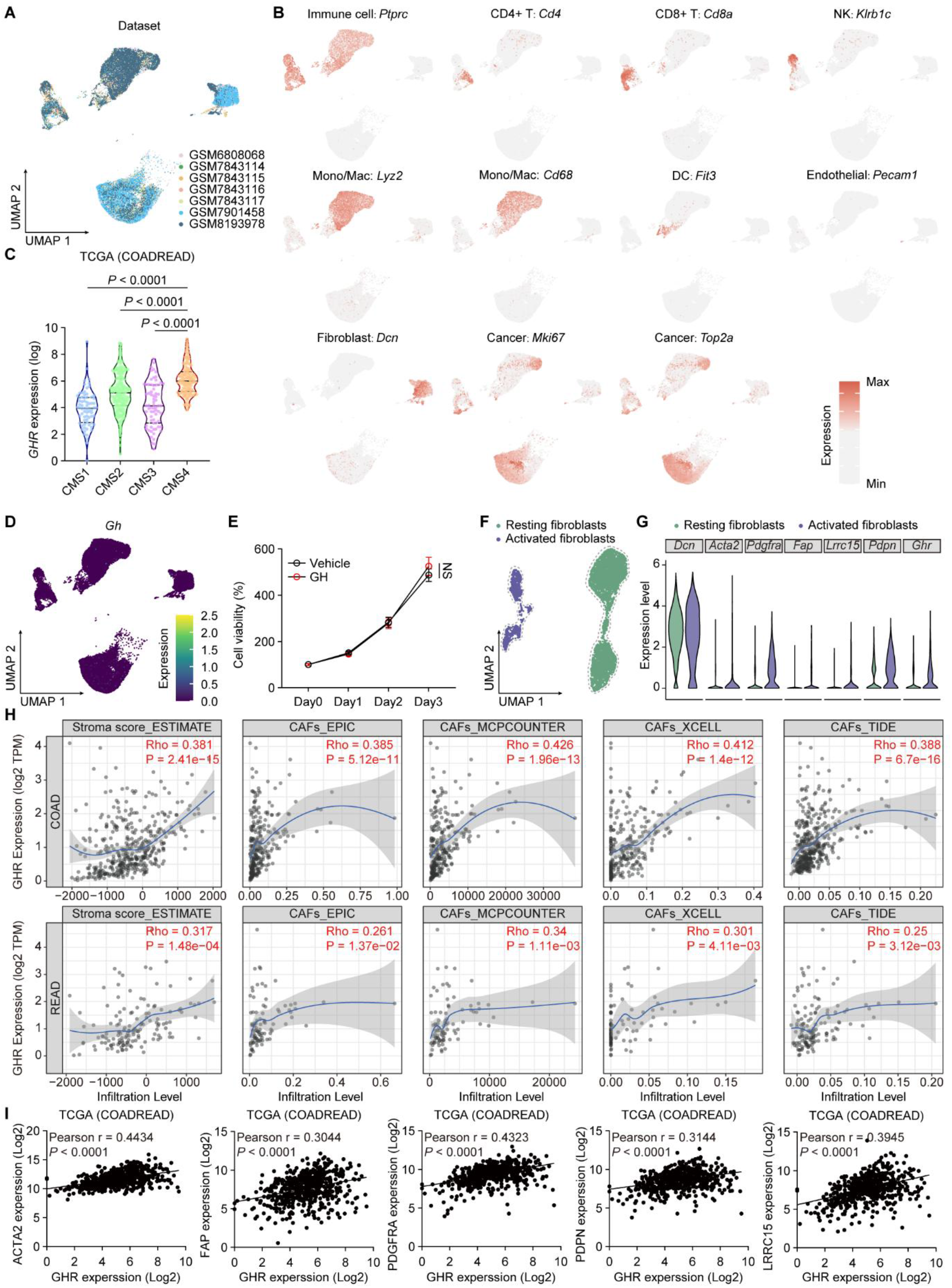
*GHR* expression is enriched in activated fibroblasts and associated with CAF signatures in CRC. **(A)** UMAP integration of seven MC38 tumor samples from four publicly available scRNA-seq datasets, colored by individual sample source. (**B**) Feature plots showing canonical marker genes for major cell populations within mouse MC38 TME, including immune cells, endothelial cells, fibroblasts and malignant epithelial cells. (**C**) Violin plots showing *GHR* expression across CMS subtypes; *n* = 71 (CMS1), 209 (CMS2), 69 (CMS3), and 135 (CMS4) patients. Solid lines, median; dotted lines, quartiles. (**D**) UMAP feature plot illustrating the cell-type-specific expression distribution of *Gh* across mouse MC38 tumor cell clusters. (**E**) *In vitro* cell viability measurement of MC38 tumor cells treated with vehicle or recombinant GH (500 ng/mL) over consecutive culture days. Cell viability was normalized to Day 0. (**F**) UMAP plot of fibroblast subpopulations stratified into resting and activated fibroblasts based on their transcriptional states. (**G**) Violin plots showing expression of *Dcn*, *Acta2*, *Pdgfra*, *Fap*, *Lrrc15*, *Pdpn*, and *Ghr* in resting and activated fibroblasts. (**H**) Correlation analysis between *GHR* expression and CAF infiltration levels in COAD and READ tissues based on TIMER3.0. Spearman’s correlation coefficients (Rho) and corresponding *P* values are annotated. (**I**) Correlation between *GHR* expression and *ACTA2*, *FAP*, *PDGFA*, *PDPN*, and *LRRC15* expression in TCGA CRC samples. Pearson correlation analysis. The data are presented as mean ± SEM. *P* values are determined by one-way ANOVA with Tukey’s multiple comparisons test (C) and two-tailed unpaired Student’s *t* test (E). NS, not significant.

**Fig. S7.**
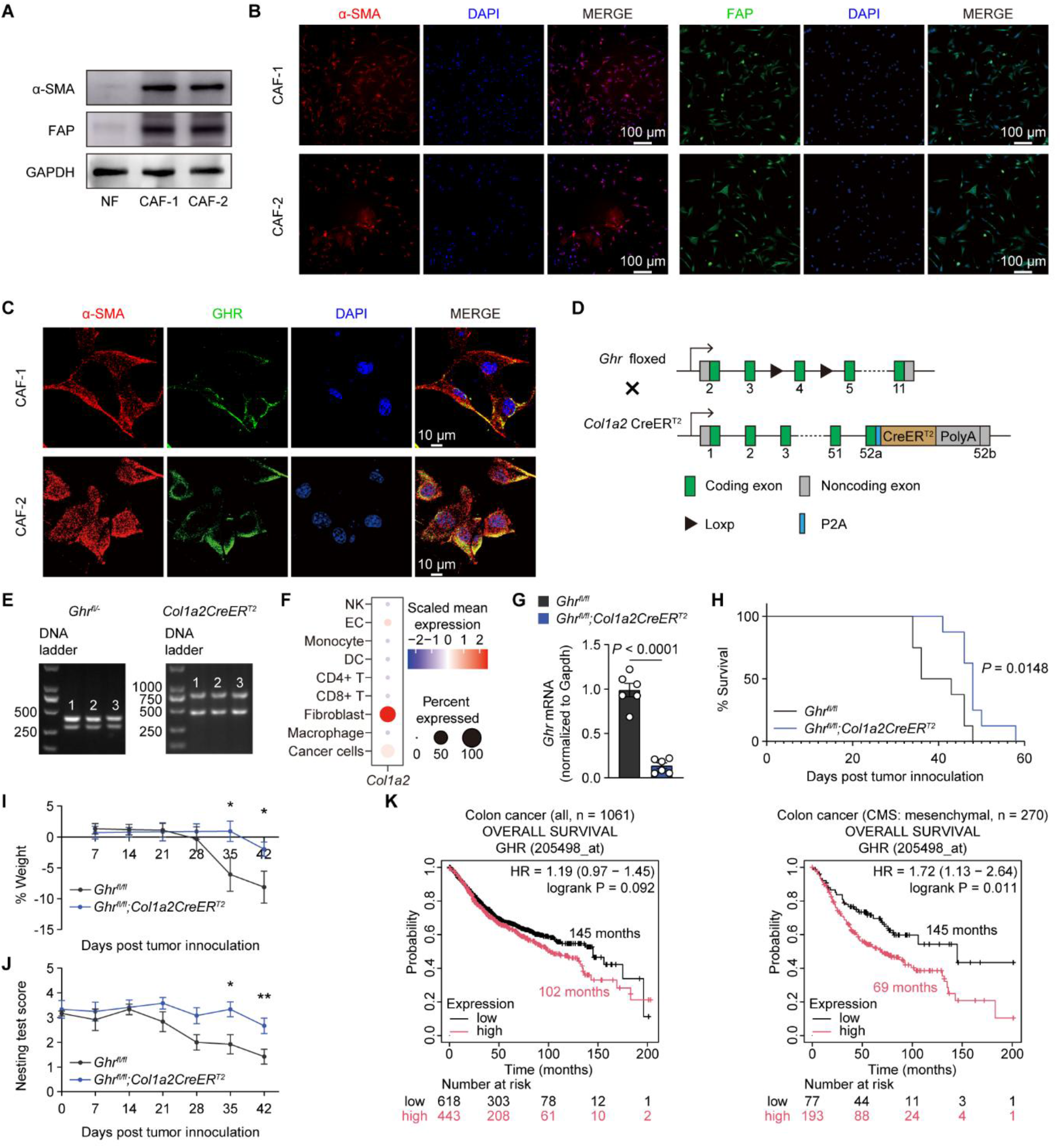
*GHR* expression in CAFs and its association with CRC progression and prognosis. (**A**) Western blot analysis to detect α-SMA and FAP expression in normal fibroblasts (NF) and two independent CAF lines, with GAPDH as the loading control. (**B**) Immunofluorescence staining of α-SMA and FAP in CAF-1 and CAF-2 cells. Scale bars, 100 μm. (**C**) Immunofluorescence co-staining of α-SMA and GHR in CAF-1 and CAF-2 cells. Scale bars, 10 μm. (**D**) Genetic strategy for the generation of *Ghr^fl/fl^;Col1a2-CreER^T2^* mouse line. (**E**) Gel verification of *Ghr^fl/fl^;Col1a2-CreER^T2^*mouse line. (**F**) Dot plot showing *Col1a2* expression across all cell subsets in mouse MC38 tumor tissues. (**G**) qRT-PCR analysis to determine the *Ghr* mRNA expression in isolated CAFs of mice carrying *Ghr^fl/fl^* or *Ghr^fl/fl^;Col1a2-CreER^T2^* after tamoxifen induction. (**H**) Kaplan–Meier survival plots of *Ghr^fl/fl^* or *Ghr^fl/fl^;Col1a2-CreER^T2^* mice that were inoculated with MC38 cells after tamoxifen injection (log-rank test). (**I** and **J**) Body weight change (I) and nesting-behavior scores (J, 0–5 scale; higher = better performance) in the same cohort as in (H). *P* = 0.0387 on Day 35, *P* = 0.0326 on Day 42 in I; *P* = 0.0245 on Day 35, *P* = 0.0036 on Day 42 in J. (**K**) Kaplan–Meier survival curves showing the prognostic value of *GHR* expression on the overall survival (OS) of all colon cancer patients (*n* = 1061, left) or CMS4 colon cancer patients (*n* = 270, right) using the Kaplan–Meier Plotter platform (*70*). The data are presented as mean ± SEM. *P* values are determined by two-tailed unpaired Student’s *t* test (G, I and J).

**Fig. S8.**
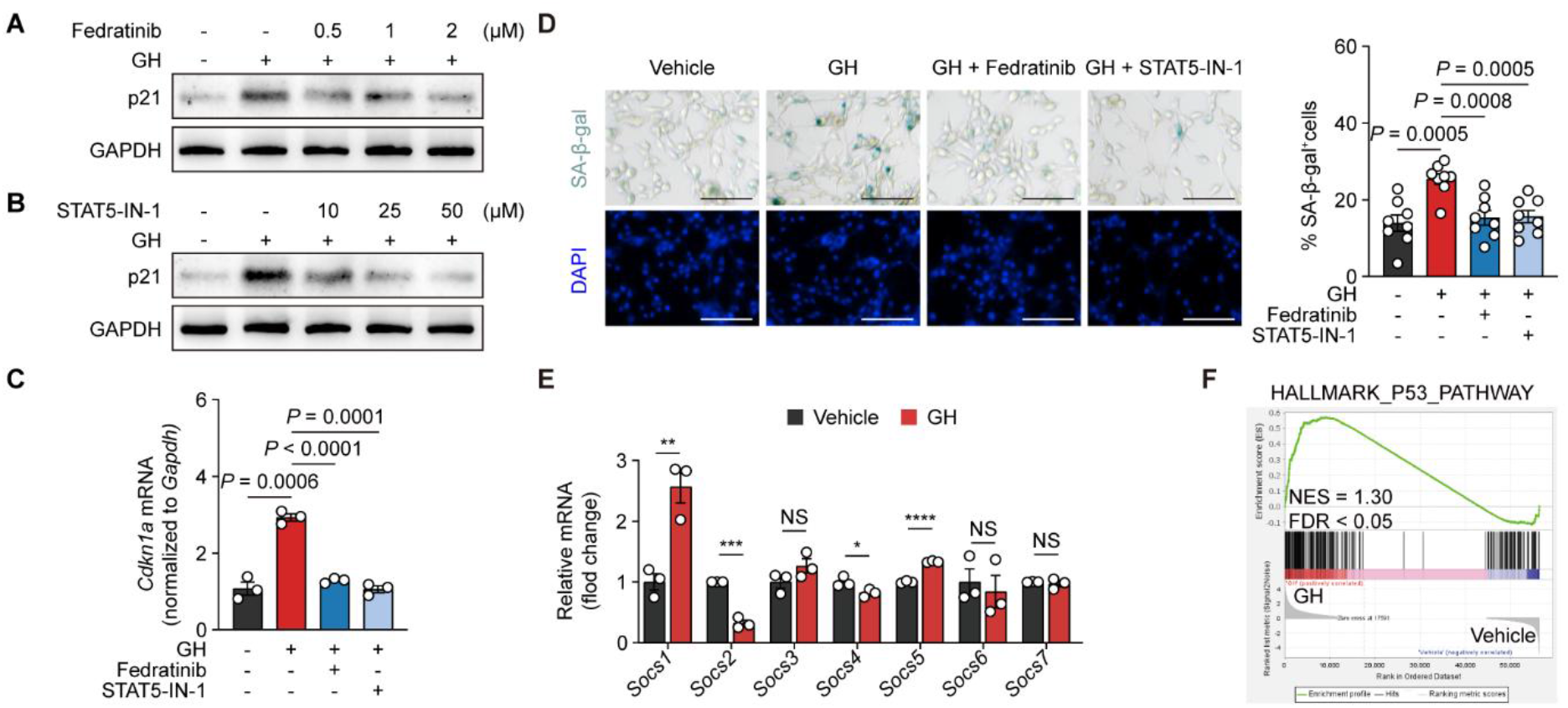
GH–GHR signaling promotes a senescence-like phenotype in CAFs through the JAK2–STAT5–p21 axis. (**A** and **B**) Immunoblot analysis of p21 in CAF-1 cells pretreated with the JAK2 inhibitor fedratinib (A) or the STAT5 inhibitor STAT5-IN-1 (B) for 30 min and then stimulated with 500 ng/mL GH for 24 h. (**C**) qRT-PCR analysis of *Cdkn1a* mRNA in CAF-1 cells pretreated with fedratinib or STAT5-IN-1 for 30 min and then stimulated with 500 ng/mL GH for 6 h (*n* = 3). (**D**) Representative images and quantitative analysis of the percentage of SA-β-gal-positive cells in CAF-1 cells pretreated with JAK2 or STAT5 inhibitor for 30 min prior to continuous GH stimulation over 7 d (*n* = 8 independent cultures). (**E** and **F**) Relative mRNA expression of *Socs* family genes (E) and GSEA enrichment plots for the p53 signaling pathway (F). RNA-seq data are derived from Fig. 5. The data are presented as mean ± SEM. *P* values are determined by one-way ANOVA with Tukey’s multiple comparisons test (C and D) and two-tailed unpaired Student’s *t* test (E).

**Fig. S9.**
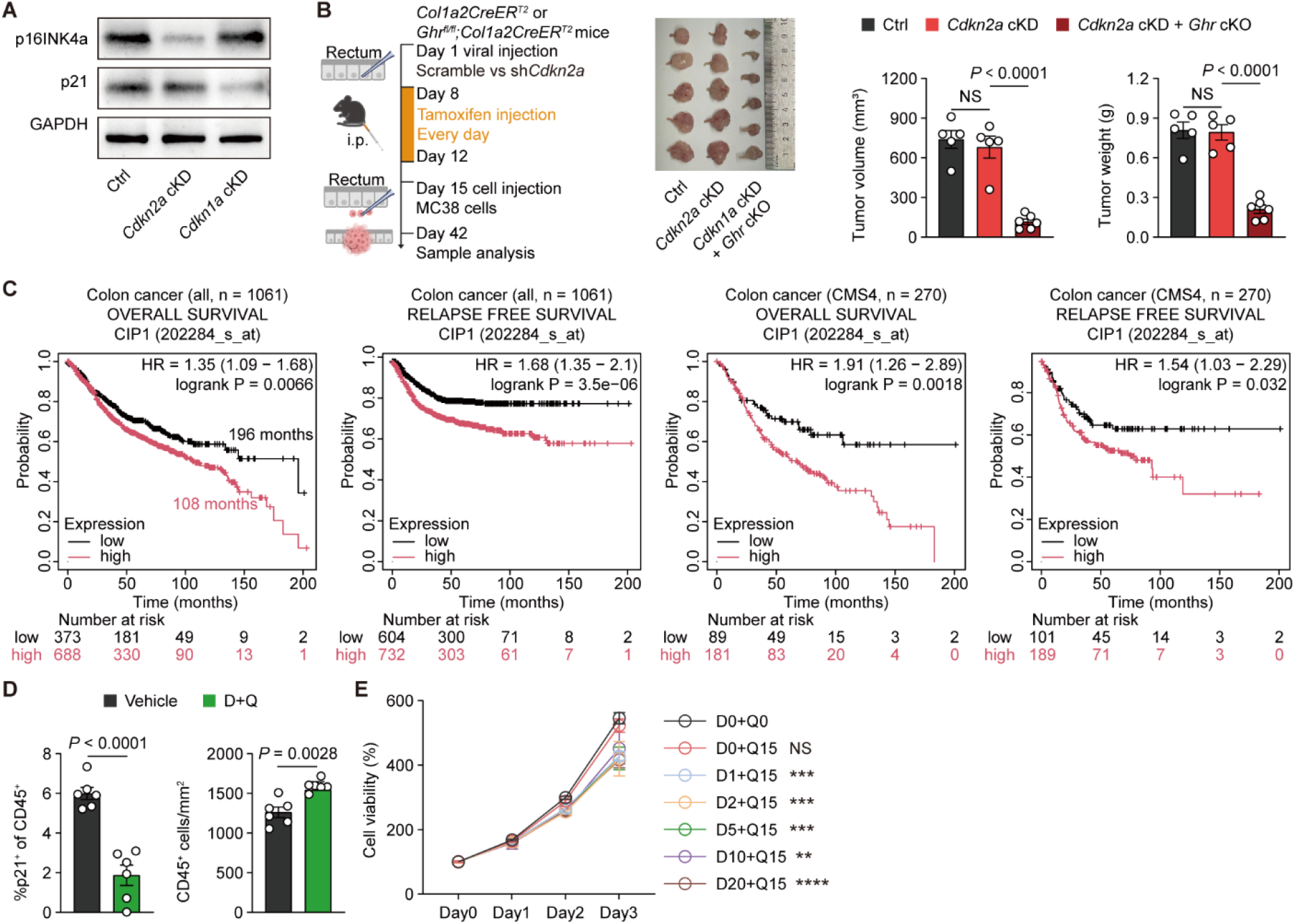
Functional analysis of senescent CAFs in CRC. (**A**) Immunoblot analysis of p16INK4a and p21 in CAFs isolated from Ctrl, *Cdkn2a* cKD, and *Cdkn1a* cKD mice. (**B**) Experimental setup (created with BioRender.com), representative rectal tumor images, and tumor quantification in Ctrl, *Cdkn2a* cKD, *and Cdkn2a* cKD *+ Ghr* cKO mice following orthotopic MC38 tumor inoculation. Tumor burden was assessed by tumor volume and tumor weight (*n* = 5–6 per group). (**C**) Kaplan–Meier analysis of the association between *CDKN1A* expression and overall survival (OS) or relapse-free survival (RFS) in all patients with colon cancer and in the CMS4 subgroup. Analyses were performed using the Kaplan–Meier Plotter (http://kmplot.com). The *P* values from the log-rank (Mantel–Cox) test, hazard ratios (HRs), and patient numbers are indicated. (**D**) Quantification of co-IF staining in MC38 tumors from vehicle- and D+Q-treated mice (*n* = 6). The left graph shows the percentage of p21^+^ cells among CD45^+^ immune cells, and the right graph shows the density of total CD45^+^ immune cells per mm^2^. (**E**) Cell viability of MC38 cells treated with vehicle (D0 + Q0), quercetin alone (D0 + Q15), or dasatinib plus quercetin (D+Q; Q fixed at 15 μM and D at 1, 2, 5, 10, or 20 μM) was assessed by CCK-8 assay at 0, 24, 48, and 72 h and normalized to the vehicle group at 0 h. Statistical comparisons at 72 h were performed against the vehicle group: D0 + Q15, *P* = 0.9567; D1 + Q15, *P* = 0.0003; D2 + Q15, *P* = 0.0001; D5 + Q15, *P* = 0.0001; D10 + Q15, *P* = 0.0058; and D20 + Q15, *P* < 0.0001. D, dasatinib; Q, quercetin. The data are presented as mean ± SEM. *P* values are determined by one-way ANOVA with Tukey’s multiple comparisons test (B and E) and two-tailed unpaired Student’s *t* test (D).

**Fig. S10.**
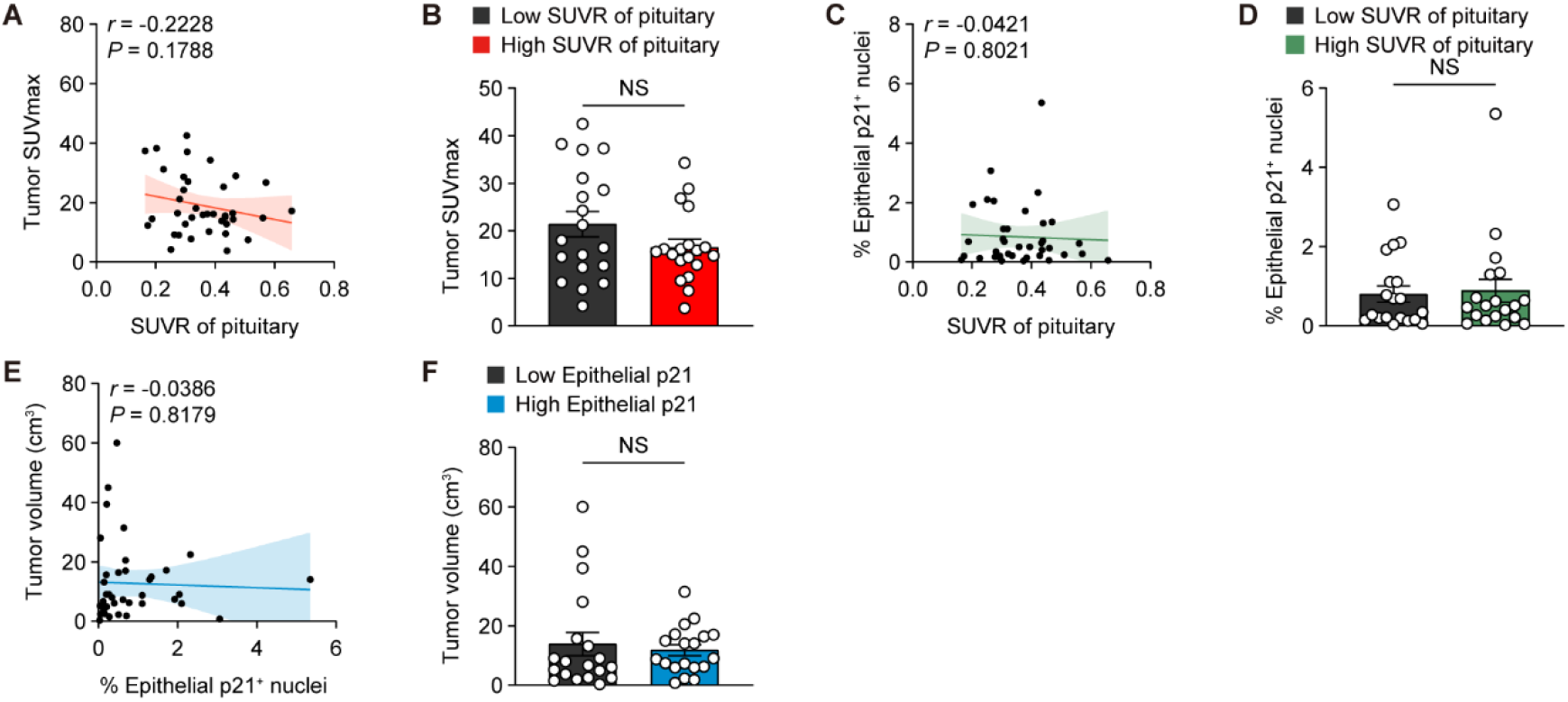
Clinical features of the pituitary–stromal axis in CRC. (**A**) Correlation analysis between pituitary ^18^F-FDG SUVR and tumor SUVmax. The solid line indicates the linear regression fit, and the shaded area represents the 95% confidence interval (*n* = 38). Pearson correlation analysis. (**B**) Comparison of tumor SUVmax between the low (*n* = 19) and high (n = 19) pituitary ^18^F-FDG SUVR groups. (**C**) Correlation analysis between pituitary ^18^F-FDG SUVR and the percentage of epithelial cells with nuclear p21 expression. The solid line indicates the linear regression fit, and the shaded area represents the 95% confidence interval (*n* = 38). Pearson correlation analysis. (**D**) Comparison of the percentage of epithelial cells with nuclear p21 expression between the low (*n* = 19) and high (*n* = 19) pituitary ^18^F-FDG SUVR groups. (**E**) Correlation analysis between the percentage of epithelial cells with nuclear p21 expression and tumor volume. The solid line indicates the linear regression fit, and the shaded area represents the 95% confidence interval (*n* = 38). Pearson correlation analysis. (**F**) Comparison of tumor volume between the low (*n* = 19) and high (*n* = 19) epithelial p21 groups. The data are presented as mean ± SEM. *P* values are determined by two-tailed unpaired Student’s *t* test (B, D and F).

